# An agent-based model of *Trypanosoma brucei* social motility to explore determinants of colony pattern formation

**DOI:** 10.1101/2025.11.04.686461

**Authors:** Andreas Kuhn, Timothy Krüger, Markus Engstler, Sabine C. Fischer

## Abstract

*In vitro* colonies of the unicellular parasite Trypanosoma brucei expand radially and establish fingering instabilities, a collective behavior known as social motility. The underlying mechanisms are thought to involve single-cell motility, chemical communication among cells, and mechanical interactions with the liquid boundary, but their relative contributions remain unclear. We aimed to determine which of the mechanisms are necessary to quantitatively reproduce the morphological characteristics of social motility. We developed a two-dimensional agent-based model that simulates colonies of 10^5^ − 10^6^ cells at single-cell resolution—two to four orders of magnitude larger than previous models. Cells are represented as point particles executing directional random walks with auto-chemotactic alignment, combined with reflective and mutable boundaries and exponential colony growth. The model was quantitatively evaluated by applying our previously established morphology metrics. We show quantitative agreement of the simulation results and experimental data in terms of colony morphology. Parameter exploration revealed that finger formation arises within a narrow range of trypanosome motility parameters that balance stochasticity and alignment, while boundary conditions modulate the speed of colony expansion. The diffusion coefficient of the chemotactic signal is the key determinant of pattern formation. Realistic behavior occurs at 2× 10^−11^ – 10^−10^ m^2^/s which corresponds to molecules of 12.1–1690 kDa. These results demonstrate that complex colony morphologies can emerge from minimal cell-level rules, suggesting testable hypotheses for the molecular drivers of trypanosome social motility. Furthermore, our approach provides a framework for dissecting the interplay between motility, signaling, and mechanical confinement in other microbial systems exhibiting collective behavior.

**Author summary:** *In vitro* colonies of the unicellular parasite Trypanosoma brucei expand radially and form finger-like patterns, a behavior known as social motility. The ability of trypanosomes to exhibit social motility has been linked to their successful journey through their insect vector, the tsetse fly, which facilitates their survival and spreading between hosts. However, the mechanisms driving this behavior are not yet clear. Experimental data suggest that the movement and growth of individual cells, their interactions with each other, and their effects on colony boundaries may all play a role. These different factors are difficult to separate in experiments. We employed mathematical modeling to investigate the relative importance of these factors and developed a model that represents individual cells. The simulations reproduced finger formation patterns similar to those observed in experiments. The results show that complex colony shapes can emerge from simple cell behaviors. We also found that the speed at which a signaling chemical spreads between cells is crucial, and that the predicted values do not match the properties of the chemicals that have been proposed so far. Identifying and testing candidate signaling molecules in experiments could be the next step. Additionally, our approach may also aid in understanding collective behaviors in other microorganisms.

## Introduction

Understanding how simple individual behaviour gives rise to complex collective phenomena is a fundamental challenge across biology. Social motility (SoMo) [1–3] in trypanosomes presents a striking example of this emergence: densely packed individual cells, swimming in an apparently random way [4] on agarose gel surfaces, form intricate colony-level patterns. Initially, these colonies exhibit a radial outgrowth and, in a second phase, produce characteristic fingering instabilities [5]. The fingers or protrusions grow in a distinct perpendicular alignment away from the colony’s centre. While the molecular and cellular mechanisms underlying individual trypanosome motility have been extensively studied [6–9], their collective behaviour is still under investigation.

Current hypotheses propose that social motility results from chemotactic responses at the colony level [10–12], with cells reacting to environmental cues such as exosomes [10], neighbouring *E. coli* colonies [11], or self-generated pH gradients resulting from glucose metabolism [12, 13]. Alternative hypotheses suggest that physical interactions, including hydrodynamic coupling, steric alignment between adjacent cells, or interactions with the liquid–solid boundary of the colony, contribute to the emergent patterns observed during collective movement [4]. The relative contributions of these chemical and physical mechanisms are unclear. Addressing this question requires an individual-based model.

Despite visual similarities between *T. brucei* colony morphologies and those observed in bacterial swarming systems [14–20], it remains unclear to what extent insights from bacterial motility can be transferred. Such bacterial systems have been modelled using continuum approaches, in which population-level behaviour is described through density fields governed by reaction–diffusion equations [16, 17, 21, 22] or by individual-based models [18, 23].

*Pseudomonas aeruginosa* stands out as the system most closely resembling *T. brucei* in both colony morphology with similar finger instabilities [24] and as well in microswimmer dynamics with similar swimming modes [25, 26] as trypoansoma brucei [27, 28]. However, a mechanistic understanding of *P. aeruginosa* swarming remains incomplete and is complicated by system-specific features, such as the secretion of rhamnolipids, which change the physical properties of the colony boundary. This limits the direct transferability of concepts to *T. brucei*.

These limitations indicate the requirement for a new individual-based modelling approach to identify the microscopic mechanisms that drive social motility in trypanosome colonies. The two-dimensional agent-based model simulates individual trypanosomes at populations of ∼ 10^5^ to 10^6^ cells, consistent with social motility assays [1, 2, 4, 5, 10]. This represents an increase of four orders of magnitude in agent numbers compared to existing models of trypanosome [27–29], which examine the swimming dynamics of individual cells, and an increase of two orders of magnitude compared to other microswimmer models [30–33] for collective behaviour.

We employ several key assumptions to make such large-scale simulations computationally feasible over the social motility timescales of up to 3 days. First, we represent each trypanosome as a point particle on a square lattice, with spacing matched to the projected area of real trypanosomes. The complex swimming motion of individual trypanosomes is reduced to a dry active matter model [34] where agents perform a directional random walk with reflective boundary conditions at the colony boundaries, capturing the essential persistent motion of these microorganisms in confined spaces [11] while abstracting away their detailed swimming dynamics [35, 36].

Second, we model the complex liquid boundary of the colony on top of the semi-liquid agarose matrix by dividing the colony space into walkable and non-walkable lattice sites. Collisions of the agents with the non-walkable lattice sites on the colony boundary make those sites progressively walkable and part of the colony. Such an approach has been successfully used before to model the growth of bacterial colonies [22, 37].

Third, we model the proposed chemical signalling and diffusion that mediates cell-cell communication by a reaction-diffusion equation and negative autochemotaxis similar to previous models for bacteria chemotaxis [16, 21, 37–40]. For this, we employ a separate, coarser grid. Our dual-grid approach balances biological realism with computational efficiency, allowing us to capture diffusion-mediated interactions while maintaining tractable simulation times.

Through analysis and comparison with experimental observations using our previously established metrics to quantify *Trypanosoma brucei* colonies [5], we demonstrate that these simplified rules enable the reproduction of the morphological characteristics of social motility. We further analyse how the individual components change the emerging behaviour and found that they exhibit distinct effects. The colony morphology is very sensitive to the motility parameters of the single cells. The parameters for boundary interactions mostly affect expansion speed. The diffusion coefficient of the chemotactic signal affects both expansion speed and colony morphology. Hence, it is a key parameter for obtaining finger-like patterns comparable to experiments. We could not identify the chemical component that most likely drives Social motility because the optimal diffusion parameter value does not match any of the previously suggested chemical components.

## Materials and methods

### Model Architecture

To model trypanosome social motility, we combined an agent-based model for the individual cells with a continuous model for chemical signalling and diffusion. The state of the model is stored in three primary data structures (Figure 1). First, a table of agents representing the trypanosomes contains for each entry a unique identifier (id), the position ***x***(*t*), and the orientation *θ*(*t*) as the angle in the 2D plane relative to the horizontal axis (Figure 1 (a)). Second, a 2D integer array called the agent-based model grid (ABM grid) models the spatial environment. Third, a 2D floating-point array called the gradient grid stores the chemical concentration values *s*(*x*). The ABM grid is a square lattice for which the lattice constant Δ*x* was set to 3.75 *µ*m, corresponding to a unit cell area of 14.06 *µ*m^2^ (see Section S.1 for more details). Each grid cell stores an integer value *k*, where:

**Fig 1.**
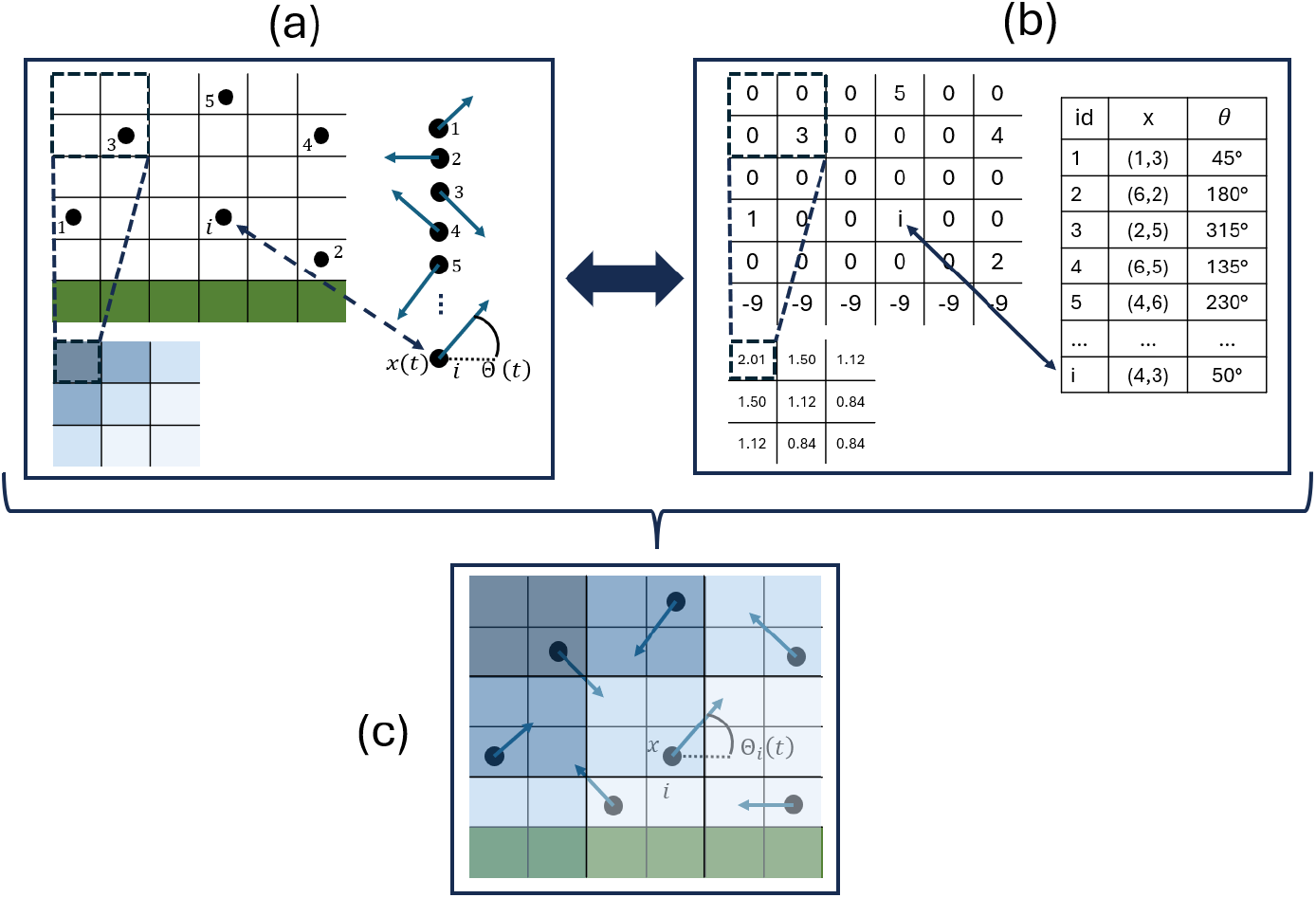
Schematic of the model architecture: (a) Conceptual representation of individual components: ABM grid (left), agent table/list (right), and gradient grid (*s*_*f*_ = 2) (below). Each agent (black dot) occupies one cell in the ABM grid; walkable cells are white, and non-walkable cells are green. The agent table stores the position *x*(*t*) and direction *θ*(*t*) (blue arrow) of all agents. With scaling factor *s*_*f*_ = 2, four ABM grid cells map to one gradient grid cell, with chemical concentration indicated by blue colour intensity. (b) Computational implementation of the data structures as in (a): The ABM grid is a 6×6 2D integer array, the agent table is a vector with position (integer tuple) and orientation (floating-point between 0° and 360°), and the gradient grid is a smaller 3×3 2D floating-point array. (c) Conceptual representation of integrated model: agents move in their current direction through varying chemical concentrations on empty ABM grid cells.

- *k >* 0 indicates the unique ID of an agent occupying that cell (only one agent per grid cell is possible),
- *k* = 0 represents an empty, walkable grid cell,
- *k <* 0 denotes a non-walkable grid cell outside the colony, with the absolute value of *k* determining the grid strength.

The gradient grid stores the local chemical concentration at each grid point and handles calculations for chemical diffusion, adsorption, and decay. This grid can be scaled down by an integer scaling factor *s*_*f*_ (e.g., 1, 2, 3, 4, …) relative to the ABM grid. Therefore, each gradient grid cell corresponds to 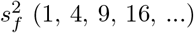 ABM grid cells.

### Initialization

The model initialisation proceeds in several sequential steps:

1. Creation of the ABM grid with desired size *L* and corresponding number of grid points.
2. Assignment of the same negative value *k*_0_ (grid strength) to all grid points.
3. Definition of the initial colony geometry by setting all grid points to zero within a circle with radius *r*_colony_ away from the center of the ABM grid.
4. Random placement of *N* agents on walkable grid points (*k* = 0) with random orientations drawn from a uniform distribution.
5. Creation of the gradient grid, scaled to the ABM grid with factor *s*_*f*_ and initialized with a small, randomly distributed chemical concentration at each grid point.

### Simulations

The simulations are performed in an iterative manner. Updates of the gradient grid, the agents, and the ABM grid are combined (Figure 2). In the following, we describe the individual parts in detail.

**Fig 2.**
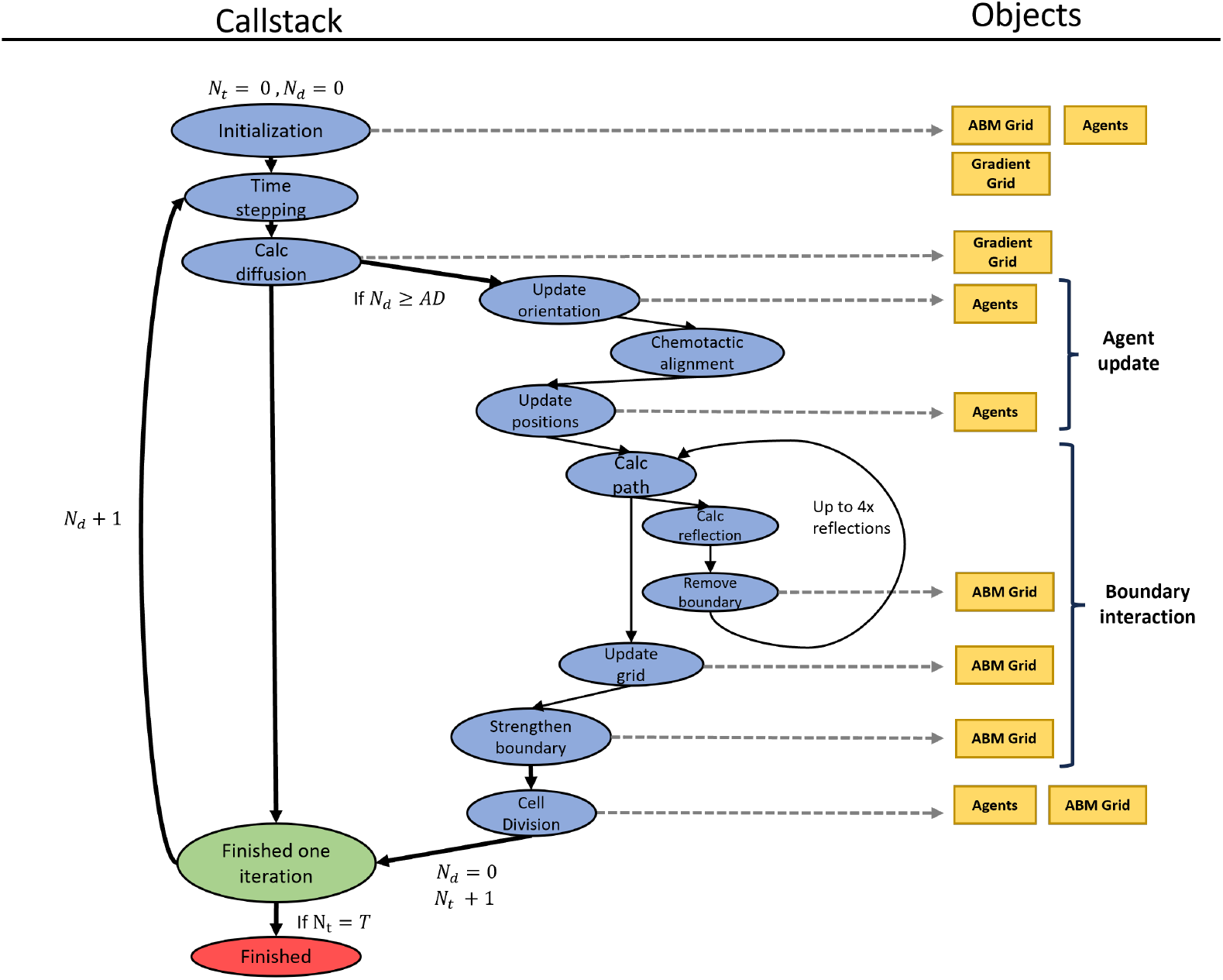
Flowchart of the internal model logic. The blue ovals represent functions that are called in consecutive order, and the yellow boxes represent the objects that they modify. The green oval represents the end of one iteration and the red oval represents the end of one model run. The model has two internal counters: *N*_*t*_, which counts the number of agent time steps Δ*t*_*a*_, and *N*_*d*_, which counts the number of diffusion time steps Δ*t*_*d*_ per Δ*t*_*a*_ to be exactly *AD*.

#### Time stepping

The model updates its components using two different time steps. The first, Δ*t*_*d*_, governs the time evolution of the gradient grid. The second time step, Δ*t*_*a*_, governs updates to agent positions, orientations, and boundary strengths in the agent-based model (ABM) grid. This separation of time scales allows for efficient simulation while maintaining the accuracy of both the chemical diffusion dynamics and agent movement. The ratio of the two time steps 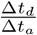 is set as *AD*. Hence, during one iteration, we perform *AD* diffusion steps and one step of updating the agents and the ABM grid (Figure 2).

#### Calculate diffusion

The model computes the time evolution of the chemical field *s*(*x, t*) on the gradient grid with a two-dimensional reaction-diffusion equation [16, 21, 37–40] with diffusion constant *D*_*s*_, decay rate *d*_*r*_ and adsorption *a*_*d*_ caused by the *N* agents at their ABM grid cells *x*_*n*_. As 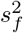 ABM grid cells are mapped to one gradient grid cell, the adsorption is normalized by that factor:

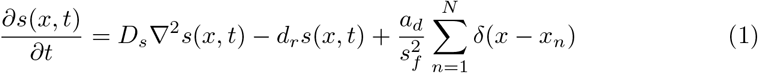

This equation is discretized and numerically solved using the finite difference method FTCS (forward time-centered space) with Dirichlet boundary conditions, implemented using the `Julia` packages `ParallelStencils` and `ImplicitGlobalGrid` [41]. This method is commonly used for solving parabolic differential equations [42–44] and was specifically chosen because its time-explicit nature allows easier integration with the agent dynamics compared to implicit methods:

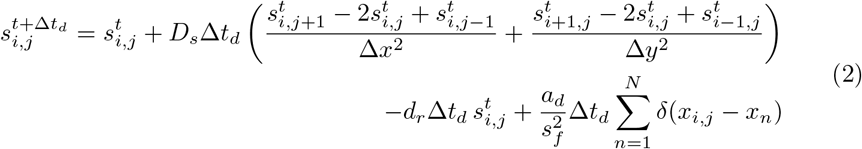

The discretized equation 2 calculates the concentration *s* at each grid cell with indices (*i, j*) at time *t* + Δ*t*_*d*_, which changes from the value of 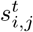 through diffusion from or to neighboring grid cells, the decay of the chemical with rate *d*_*r*_, and the adsorption of the chemical with rate *a*_*d*_. The Dirac delta function (*δ*()) is only non-zero when *x*_*n*_ is one of the 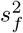 ABM grid cells mapping to the gradient grid cell with position *x*_*i,j*_ (Fig. 1).

The maximum time step size Δ*t*_*d*_ to ensure numerical stability can be derived using von Neumann stability analysis [44–46] (see Section S.2 for more details):

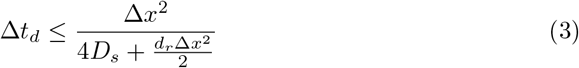

#### Agent update

In every time step Δ*t*_*a*_, each agent updates its orientation and position. First, the orientation is updated according to:

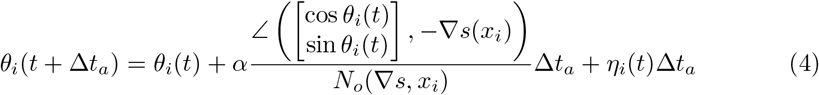

where the angle between two vectors is calculated as:

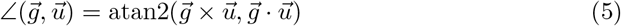

and the normalization function *N*_*o*_ is:

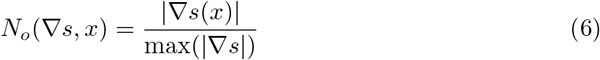

The agents try to turn their current direction towards the negative gradient of the chemical field *s*(*x*_*i*_) at their position *x*_*i*_. The speed of this negative chemotactic alignment process is determined by the maximum gradient in the system through the normalization factor *N*_*o*_ and a tunable turning rate parameter *α*, creating a relative response: For the case where *α* = 1, an agent at the location of the steepest gradient in the system will completely align its direction in one time step, while agents in regions with weaker gradients will align proportionally slower. When *α <* 1, the alignment speed is linearly scaled down by this factor. The term *η*_*i*_(*t*) models the stochastic component of the agent’s movement and alignment process and is drawn from a normal distribution with mean 0^*°*^ and standard deviation *η*.

Each agent’s position is updated according to:

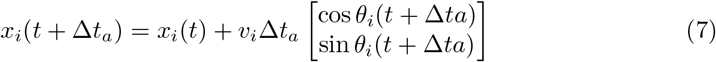

The agents move from their current position *x*_*i*_(*t*) in the direction *θ*_*i*_(*t* + Δ*t*_*a*_) with velocity *v*_*i*_, which is drawn from a normal distribution with mean *v*_0_ and standard deviation *v*_*σ*_. Since *x*(*t* + Δ*t*_*a*_) is a vector of two continuous values, it is rounded to the nearest discrete grid cell on the ABM grid. Such an update scheme would be labeled as microscopic dry active matter [34], or in other terminology, modified active Brownian particles with auto-chemotactic alignment.

Agent update depends on the time step Δ*t*_*a*_. Its value must strike a balance between two opposing constraints. If Δ*t*_*a*_ is too small, the time discretization becomes overly fine, potentially introducing discretization artefacts: agents may move only one grid cell, or remain stationary, per step, which distorts movement dynamics [47]. Conversely, if Δ*t*_*a*_ is too large, agents may traverse unrealistically long distances in a single step. This results in other discretization artefacts, such as unnaturally straight trajectories, which contradict experimental observations of cell motility [4]. In addition, rapid changes in the local environment, such as the number and identity of neighbouring agents or proximity to boundaries, impair biologically plausible agent-agent interactions.

To balance these constraints, Δ*t*_*a*_ was chosen to allow agents to move on average three ABM grid cells per time step in their current direction. This corresponds to a distance of 11.25 *µ*m, which is shorter than the typical cell length of a procyclic trypanosome (∼ 15–25 *µ*m [4, 48]), thereby preserving local agent-agent interactions.

For typical model parameters, the ratio 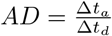 ranges between 1 and 300. If necessary, the diffusion time step Δ*t*_*d*_ is slightly reduced from its maximum allowable value to ensure that *AD* is always an integer. This guarantees a consistent number of diffusion time steps Δ*t*_*d*_ per agent time step Δ*t*_*a*_ throughout the simulation.

#### Boundary Interactions

The model simulates agent interactions with the colony boundary through reflective boundary conditions, whereby agents are reflected upon encountering a non-walkable grid cell on their path. Whilst real trypanosomes do not undergo physical reflection from colony boundaries, they typically remain at boundaries for only brief periods (a few seconds) before reorienting [4]. Therefore, reflective boundary conditions represent a justified simplification that captures the average behaviour at the agent numbers simulated here.

The lattice constant Δ*x* and time step Δ*t*_*a*_ are set to values so that agents move an average of three grid cells per time step. Since their movement paths can therefore interfere with non-walkable grid cells with *k <* 0, their movement paths are divided into segments of single grid cells. At each segment, the model checks for the presence of non-walkable grid cells with *k <* 0. If one is encountered, the agent is reflected.

The reflection process approximates the local boundary morphology by constructing a circle with radius *R*_*t*_ around the collision point (the first grid cell in an agent path segment with *k <* 0). The surface tangent is approximated using the two most distant non-boundary points neighbouring a boundary point. The agent is then reflected off this tangent such that the incoming angle *β*_1_ equals the outgoing angle *β*_2_ (Fig. 3).

**Fig 3.**
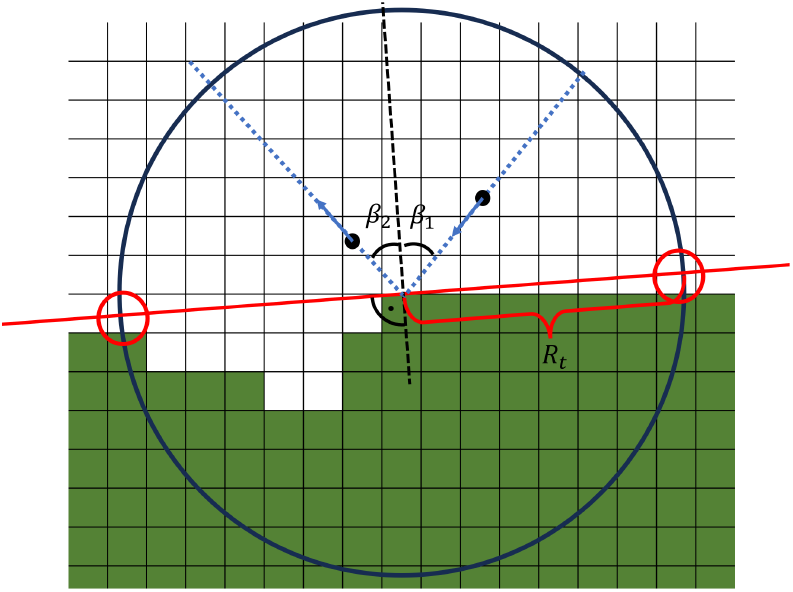
Schematic representation of the boundary reflection mechanism. The boundary morphology is approximated at the collision point with a circle of radius *R*_*t*_, where its most distant surface points form a tangent. The agent is reflected with the outgoing angle *β*_2_ equal to the incoming angle *β*_1_ relative to the tangent.

If an agent collides with a non-walkable grid cell on the colony boundary, the boundary weakens (Fig. 4). This process is modelled through a change in the value *k*, which is initialized at all non-walkable grid cells with a negative number *k*_0_ representing the grid strength. Upon collision, all grid cells with *k <* 0 within a circle of radius *R*_*c*_ (distinct from *R*_*t*_) centred on the colliding agent have their *k* values increased by 1. To address lattice anisotropy effects, not a perfect circle with radius *R*_*c*_ was used, but rather a hybrid shape combining a circle and a diamond (see Section S.3 for more details). If *k* reaches zero, the non-walkable cells become walkable space and are removed from the boundary. Similar approaches have been successfully used to model bacterial colony growth [22, 37].

**Fig 4.**
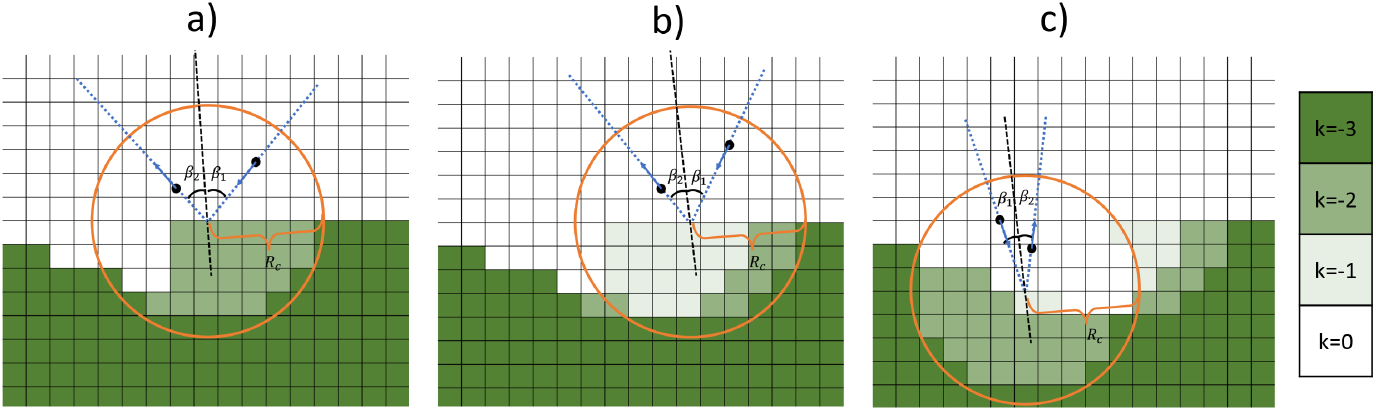
Schematic representation of the boundary removal mechanism for an initial boundary strength *k* = − 3 and three subsequent agent collisions (a, b, c). Upon agent collision with a boundary, the boundary strength *k* increases by 1 in all boundary cells within radius *R*_*c*_. When *k* reaches zero after multiple collisions, boundary cells are converted to walkable space.

Due to potentially complex local boundary morphology forming structures such as narrow channels, this collision-reflection-weakening process can occur up to 4 times in a single time step (Fig. 2). Such complex morphology can also create cases where no unambiguous tangent can be determined; in these situations, a fallback mechanism is invoked where the agent is not reflected but simply reverses its original direction.

To simulate a simple version of surface tension, a grid recovery rate *g*_*rc*_ is implemented. This parameter decreases the absolute value of *k* in every boundary grid cell by a random integer drawn from a Poisson distribution with *λ* = *g*_*rc*_ · Δ*t*_*a*_ in each time step Δ*t*_*a*_ until it reaches its initialization value *k*_0_, effectively strengthening the boundary over time if it is not further weakened by agent collisions.

#### Cell Division

Cell growth is modelled as exponential, based on population doubling times *τ*_*d*_ observed in experiments [3, 4, 12]. From the doubling time, we derive the growth rate *r* as:

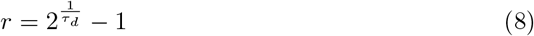

In each time step Δ*t*_*a*_, each agent has a probability *r* · Δ*t*_*a*_ of dividing and forming a new agent in a neighbouring empty grid cell. If no neighbouring empty grid cell is available, the agent cannot divide due to spatial constraints. The new agent is assigned a random orientation independent of the parent agent. Throughout the results and discussion sections, we primarily use the doubling time *τ*_*d*_ rather than the growth rate *r* to describe growth dynamics, as it provides a more intuitive interpretation of population expansion.

### Model visualisation

The visualisation integrates all three components of the model into a single composite image (Figure 5). The agents are represented by a vector plot, where each arrow corresponds to one agent. These arrows indicate the current direction of movement, with additional colour-coding to distinguish agent orientations.

**Fig 5.**
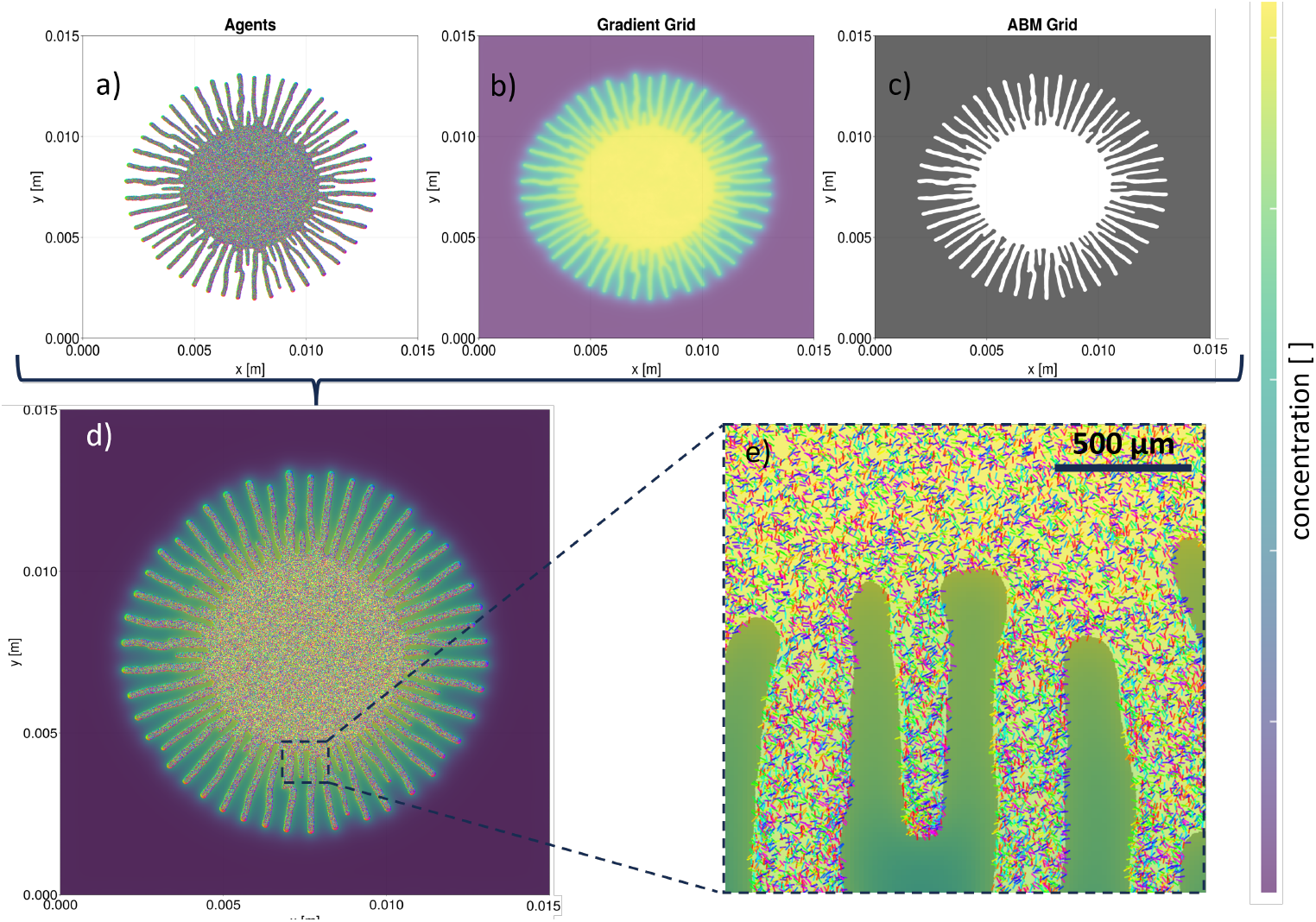
Visualisation scheme of the model for default parameters (Table 1) with *L* = *x* = *y* = 15 mm and after 9 hours: (a) The ∼ 5 *×* 10^5^ agents are displayed as a vector plot where each agent is represented by an arrow pointing in its current direction. The direction is additionally colour-coded. Due to the number of agents, it is not possible to distinguish individual agents. (b) The gradient grid is visualized by a heat map where relative concentration values map to colours (colourbar on the right). (c) The ABM grid is displayed as a binary mask where non-colony regions (*k <* 0) are shaded gray and regions within the colony (*k* ≥ 0) are transparent. (d) The three components of the model are plotted together in one single plot containing all information. (e) Zoomed-in view of the complete plot to better visualize individual agents in their crowded environment. The agents are colour-coded according to their direction.

The chemical gradient is displayed as a heat map with concentration values mapped to a colour scale. Since it is not yet clear which chemical is responsible for chemotactic alignment in trypanosome colonies and concentrations for the candidate chemicals have not been measured either, the absolute values of the concentration are less important than the relative concentration gradients in the system, which drive agent behaviour. Therefore, in all visualisations, the colourbar is auto-scaled to the maximum concentration present in each specific image. This approach allows us to visualize the relative concentration gradients consistently across all images using the same colour scaling, even though the absolute concentration values may differ substantially between different simulation conditions or time points.

The ABM grid is visualized as a binary mask where non-colony regions appear gray while colony regions are transparent. These three visual elements are layered to provide a comprehensive view of the entire system, enabling intuitive observation of emerging patterns during simulation.

Most images shown in the subsequent chapters do not display individual agents, as it is not possible to discern individual agents for simulations with realistic agent numbers, and most analysis in the following sections focuses primarily on the morphology of the ABM grid.

### Implementation

The code for implementing the model and the simulations was designed with modularity and expandability as primary considerations. While this approach introduces some computational overhead, as some code parts are not optimized for maximum memory efficiency, it offers significant advantages for future development. The data structures have been chosen to allow a straightforward integration of additional model parameters or agent attributes in subsequent model iterations.

The code is written in the programming language `Julia` [49], which aligns with the model’s design philosophy. `Julia` combines the readable, accessible syntax of high-level languages with performance comparable to traditional compiled languages like `C` and `Fortran`. This combination is particularly valuable for scientific computing applications where accessibility and runtime performance are critical. `Julia’`s growing ecosystem of scientific libraries also provides valuable tools for efficient numerical computations, particularly for the reaction-diffusion equation.

The code is available at https://github.com/scfischer/2026-Kuhn-et-al and will be archived at Zenodo upon publication.

### Simulation Modes

The simulations can be run in two distinct modes:

The first is the interactive mode, where only the current state of the simulation is stored in memory and the output is displayed with live animation using the **Makie.jl** plotting package [50]. In this mode, the parameter values can be dynamically adjusted during the simulation run to study their influence on model behaviour. The interactive mode can simulate up to 10^6^ agents on modern CPUs (2025) in real-time, meaning one second in the model corresponds to one second or less in real time. The computation times of the model scale almost linearly with the number of agents and grid cells used.

The second is the non-interactive mode, where the simulation runs with predetermined parameter values without graphical output. Here, the state of the simulation is saved at specified time points for later analyses. This mode is approximately twice as fast as the interactive mode and is primarily used for parameter sweeps in multiple parallel instances on workstations or clusters with sufficient memory per core. The saved data are subsequently used for analyses and visualisation.

### Parameter Space

The model possesses a large parameter space, the parameters of which can be classified into four distinct categories:

a. Parameters corresponding to measurable quantities that have been experimentally determined (e.g., growth rate *r* of trypanosomes in social motility assays). These experimentally measured values were adopted as default values in the model.
b. Parameters corresponding to measurable quantities that have not yet been measured (e.g., adsorption rate *a*_*d*_, decay rate *d*_*r*_, and diffusion constant *D*_*s*_ of the proposed chemotactic substance [11, 13]). The reasons these remain unmeasured vary, but generally stem from either measurement difficulties or a lack of prior scientific motivation to quantify them.
c. Parameters that do not directly correspond to measurable physical quantities (e.g., boundary strength *k* and grid recovery rate *g*_*rc*_). These parameters arise from simplifications in the model. For example, the real colony boundary represents a complex hydrodynamic interface between a semi-fluid (agarose) and a fluid medium (nutrition solution containing trypanosomes), which can exchange liquid while remaining separated by surface tension. Although the properties of this interface could theoretically be measured, incorporating them would require modelling the interface with comparable complexity.
d. The discretisation of time and space is also a model parameter; the values chosen/calculated for these are explained above and in Section S.1.

For parameters in categories (b) and (c), default values were determined through extensive testing, primarily in interactive mode. These values were selected to produce colony morphology dynamics similar to those observed and quantified in experiments [5]. A detailed analysis of their influence is shown in the results section.

All parameters are shown in Table 1 together with their category and default values. There are two values given for the doubling time *τ*_*d*_ as different experiments reported either doubling times of 9 h [3, 12] or 20 h [4]. We tested both values.

**Table 1.** Model parameters categorised by type with their default values and units.

| Parameter | Description | Type | Default Value | Units | Source |
| --- | --- | --- | --- | --- | --- |
| $\tau_d$ | Doubling time | a | 9.0/20.0 | $h$ | [3, 4, 12] |
| $v_0$ | Mean agent velocity | a | 5 | $\mu\text{m/s}$ | [4] |
| $v_\sigma$ | Velocity stdd | a | 2.5 | $\mu\text{m/s}$ | [4] |
| $D_s$ | Diffusion coefficient | b | $7 * 10^{-11}$ | $\frac{m}{s^2}$ | - |
| $a_d$ | Adsorption rate | b | 0.000001 | $s^{-1}$ | - |
| $d_r$ | Decay rate | b | 0.001 | $s^{-1}$ | - |
| $k_0$ | Boundary strength | c | -8000 | - | - |
| $g_{rc}$ | Grid recovery rate | c | 7 | $s^{-1}$ | - |
| $R_c$ | Collision radius | c | 16 | grid cells | - |
| $R_t$ | Tangent approx r | c | 15 | grid cells | - |
| $\eta$ | Orientation noise | c | 0.2 | rad/s | - |
| $\alpha$ | Turning rate | c | 0.1 | rad/s | - |
| $s_f$ | Grid scaling factor | d | 4 | - | - |
| $\Delta x$ | ABM lattice constant | d | 3.75 | $\mu\text{m}$ | - |
| $\Delta t_a$ | Agent time step | d | 2.25 | s | - |
| $\Delta t_{\text{diff}}$ | Diffusion time step | d | 0.75 | s | - |

The time steps Δ*t*_*a*_ and Δ*t*_*d*_ are dynamically calculated quantities that change depending on the diffusion coefficient *D*_*s*_, the mean agent velocity *v*_0_, the decay rate *d*_*r*_, and the lattice constant Δ*x*.

Unless otherwise specified, all presented data are from simulated colonies starting with *N* = 2.5× 10^5^ agents in a circular geometry with a diameter of 3 mm. This represents a similar trypanosome density to that observed in the well-quantified experiments from Kruger et al. [4, 5], which serve as a blueprint for our model. In these experiments, approximately 10^6^ agents are situated in colonies with diameters of 6 mm. Initial testing showed that this downscaling of the system by a factor of four did not change the overall behaviour but reduced computation times by a factor of approximately eight. This computational advantage aligns with other experimental observations, which demonstrated unchanged social motility activity in smaller colonies of *N* = 2×10^5^ trypanosomes [12], hence its adoption in our simulations.

Another performance-relevant quantity to determine before each simulation run is the edge length *L* of the simulated space. The computation time for the diffusion equation on the gradient grid scales with *L*^2^. Since we use Dirichlet boundary conditions, the colony cannot expand beyond the grid boundaries. Therefore, *L* should be large enough to prevent the colony from reaching the boundaries during the simulation period, yet as small as possible to maintain reasonable computation times. After testing, we determined that *L* = 15 mm for simulations running 10 h and *L* = 30 mm for simulations running 20 h satisfied these conditions well, and we used these values throughout our analyses.

One important observation from our testing is that the variation between model runs with identical parameters is remarkably small. Given this high reproducibility and the significant computational demands, we opted to run each parameter set only once for the simulations presented below, thereby substantially reducing overall computation time without compromising the validity of our conclusions.

### Analysis

The simulation results were analyzed by characterizing the ABM grid. We measured the area *A* of the colony at each time point as the number of colony grid points and normalised the increase in area with respect to the initial time point. Hence, for a given time point *t*^∗^ and initial time point *t*_0_, we calculated the relative area increase of the simulated colony as 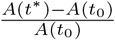. Furthermore, we counted the number of agents at each time point to obtain the cell number and calculated the sum of adsorbed material on the gradient grid. The cell density was calculated as 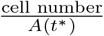. To quantify the colony morphology, we used surface roughness *W*, coefficient of variation *CV*, relative maximum peak height *MaxP*1 and mean Fourier amplitude *I*_*k*_. These morphological metrics are based on an angular metric and a pair correlation metric and were previously introduced for trypanosome colonies [5].

## Results

### Simulated colonies show two-phase expansion behaviour

To establish baseline behaviour and validate our model against experimental observations, we simulated the system for 20 h using default parameter values (Table 1) with two biologically relevant doubling times: *τ*_*d*_ = 9 h (Fig. 6) and *τ*_*d*_ = 20 h (Fig. 7), corresponding to different experimental conditions [3, 4, 12].

**Fig 6.**
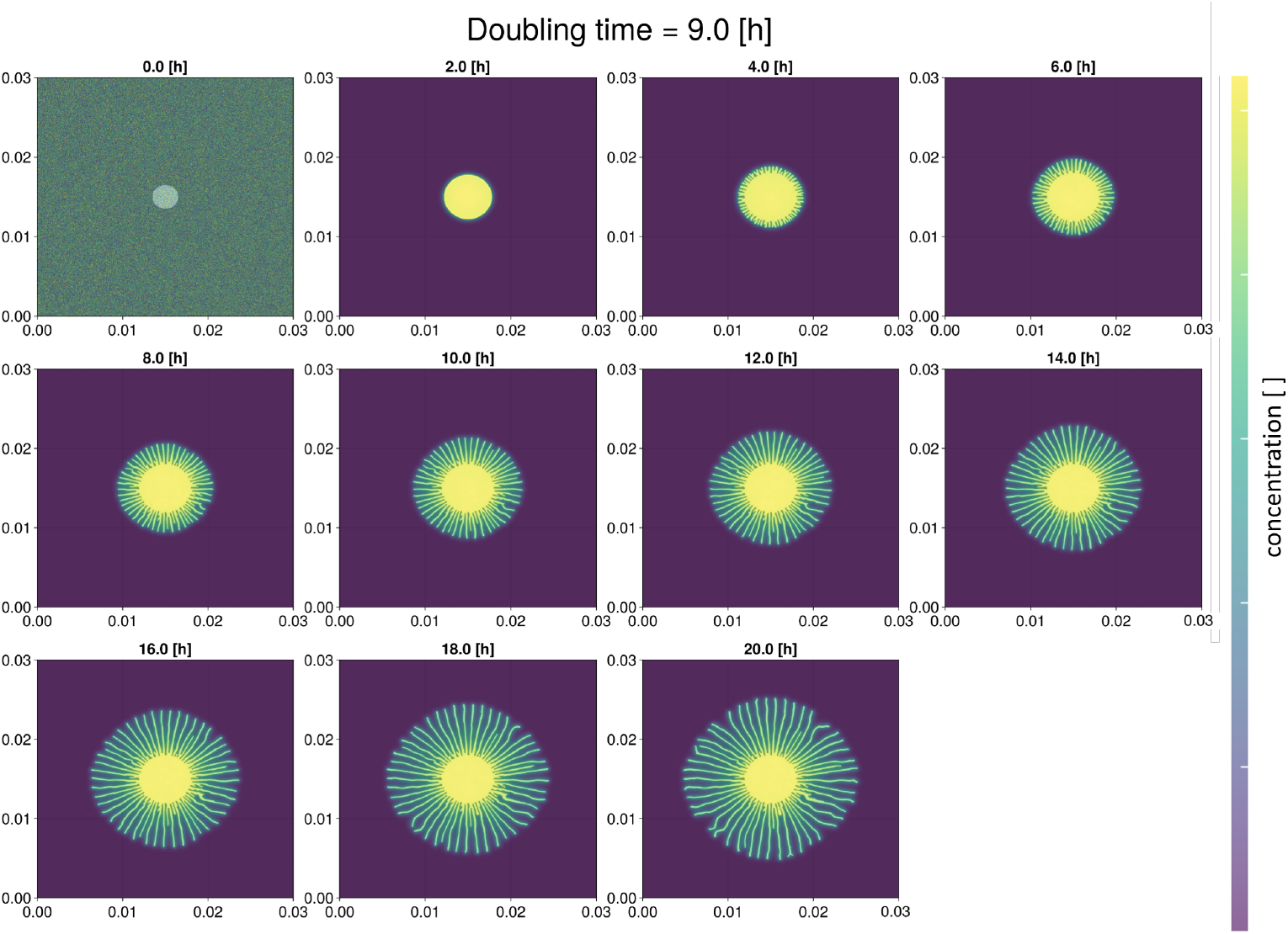
Time evolution of the colony morphology over 20 hours (2-hour intervals) with *τ*_*d*_ = 9 h and default parameters (Table 1). Initial noise in the gradient grid reflects random initialization. For details of visualisation see Materials and Methods.

**Fig 7.**
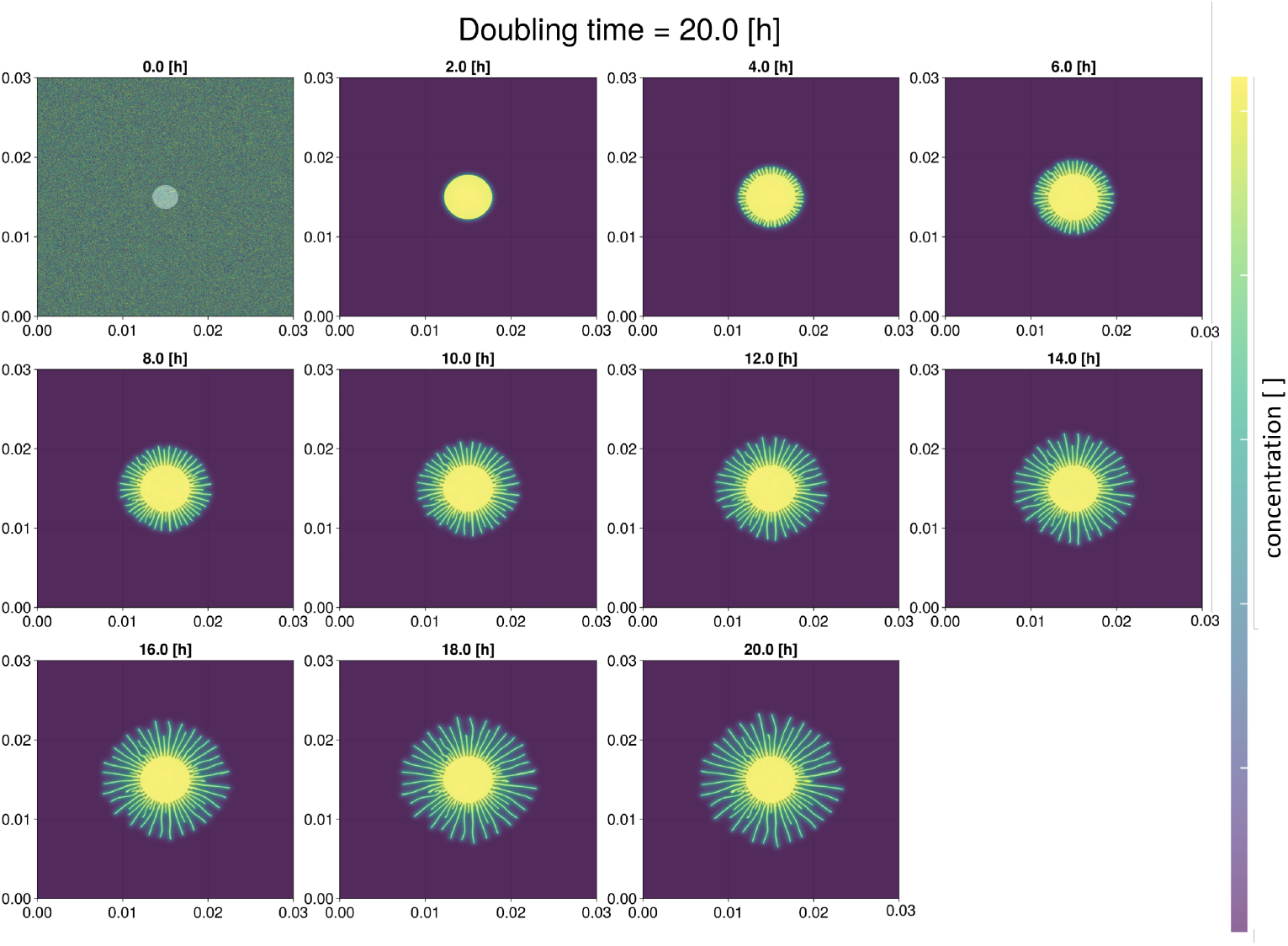
Time evolution of the colony morphology over 20 hours (2-hour intervals) with *τ*_*d*_ = 20 h and default parameters (Table 1). Initial noise in the gradient grid reflects random initialization. For details of visualisation see Materials and Methods.

Both simulations exhibit a two-phase expansion: initial circular growth (0-2 h) followed by finger-like expansion. The faster-growing colony (*τ*_*d*_ = 9 h) develops approximately 50 fingers with more uniform morphology, while the slower case (*τ*_*d*_ = 20 h) produces approximately 45 fingers with greater variability in length.

During the circular phase (0–2 h), the relative area increase is nearly identical across colonies, despite differences in doubling time (Fig. 8). The number of cells increases exponentially throughout this phase. Decay and adsorption balance within the first hour. Afterwards, the amount of adsorbed material increases exponentially, mirroring cell growth. Cell density reaches a minimum during the first 10 h, coinciding with the morphological transition. All measures increase more rapidly in the faster-growing colony.

**Fig 8.**
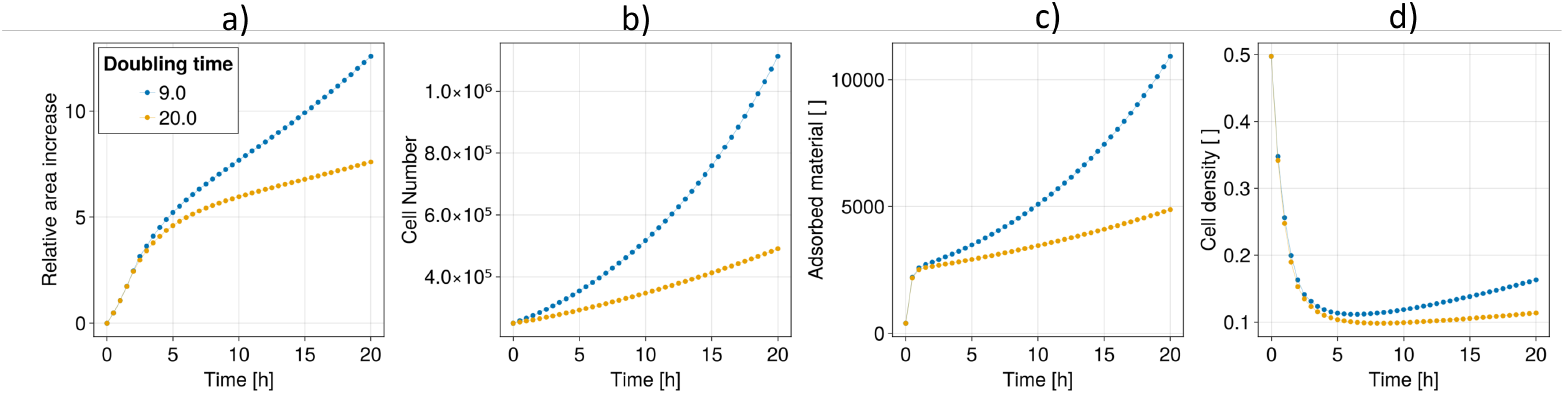
Temporal evolution of colony properties in 30-minute intervals for two doubling times and default parameters (Table 1). The results are for one simulation run each due to very low variability between runs (see Materials and Methods for details). (a) Relative area increase, (b) Cell number, (c) Adsorbed material, (d) Cell density. Note the density minimum at 4 h, coinciding with the morphological transition.

We employed our established morphological metrics that are able to quantify the change from circular to finger-like growth by asserting values of zero to a colony growing perfectly circular and increasing to higher values for more finger-like growth (Fig. 9, [5]).

**Fig 9.**
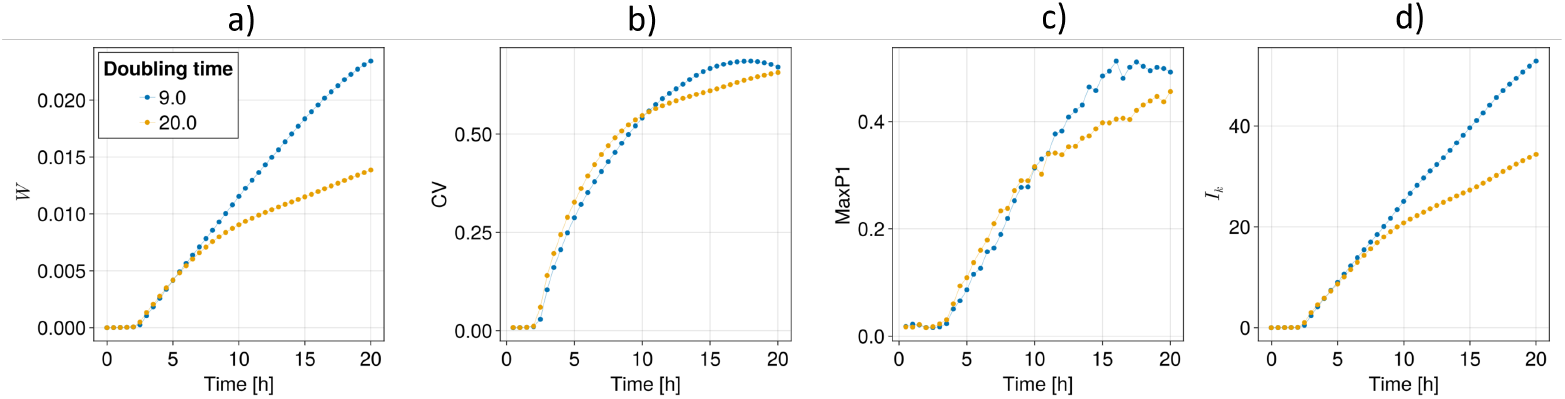
Temporal evolution of morphological metrics in 30-minute intervals for two doubling times and default parameters (Table 1). The results are for one simulation run each due to very low variability between runs (see Materials and Methods for details). (a) Surface roughness *W*, (b) Coefficient of variation *CV*, (c) Relative maximum peak height *MaxP*1, and (d) Mean Fourier amplitude *I*_*k*_.

All metrics clearly detect the initial circular growth phase, showing values at or near zero during the first two hours of the simulations. Subsequently, all metrics capture the transition from circular to finger-like growth as their values begin to increase. The coefficient of variation of the angular metric (*CV*) shows the highest sensitivity to this transition as it already increases at 1.5 h.

The roughness (*W*) and the mean Fourier amplitude (*I*_*k*_) exhibit patterns of steady increase for both doubling times. In contrast, the *CV* and *MaxP*1 metrics increase more slowly after 10 h. For *τ*_*d*_ = 9 h, they eventually saturate. Both quantities particularly represent expansion that is perpendicular to the center point of the colony at *t* = 0 h [5]. After 10 h, this becomes less pronounced because single fingers begin to diverge from the initial straight paths and create a less uniform expansion front (Fig. 6). In summary, the model exhibits the two-phase expansion behaviour observed *in vitro*. Our established metrics for *in vitro* Trypanosoma colonies successfully quantify the behaviour of the *in silico* colonies [5].

### Simulations can reproduce experimental morphological metrics

We compared our results with experimental data from [4] and [5]. Visual comparison shows similar morphologies but with more fingers in the simulated data compared to the experimental data. Additionally, our simulations show the perpendicular alignment of trypanosomes at the colony boundary that has been observed experimentally (Fig 5, [4]).

We further conducted a quantitative comparison of the simulation results with respect to the experimental data for two different conditions [5]. In experiment 1, 10^6^ cells were seeded. The colonies showed a relative increase in area of 1 after 20 h and developed 15 fingers. In experiment 2, twice the number of cells was seeded. The colonies showed a relative increase in area of 3.5 after 20 h and developed 30 fingers. Compared to this experimental data, the simulated colonies grow significantly faster, with a relative area increase of 7.5 for a doubling time of 20 h and 13 for a doubling time of 9 h, respectively (Fig. 8a).

To enable a quantitative comparison of the morphological metrics despite the differences in relative area increase, we plotted the simulation results and the experimental data against relative area increase (Fig. 10 and 11). The simulated colony with *τ*_*d*_ = 20 h transitions to finger-like growth at a smaller relative area increase compared to the simulated colony with *τ*_*d*_ = 9 h (Fig. 10), indicating that increased growth rate not only accelerates colony expansion but also changes the nature of that expansion. For the experimental data, experiment 1 exhibits clearer fingering at a smaller relative area increase when compared to experiment 2 (Fig. 11). We find that for experiment 2, the orders of magnitude of the simulation results agree well with the experimental data. For experiment 1, the morphological metrics are larger than for the simulations.

**Fig 10.**
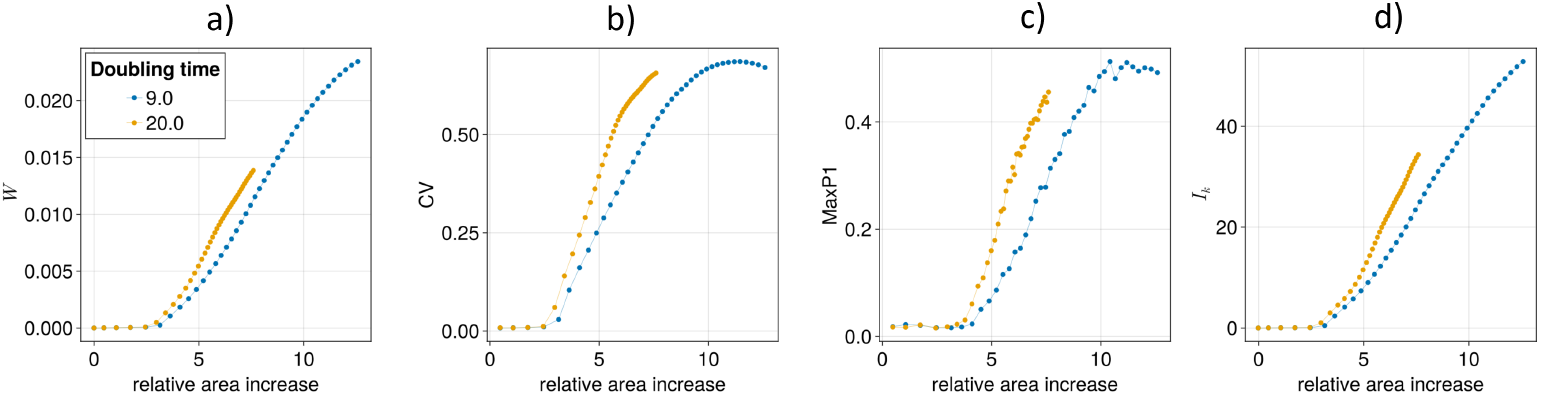
Evolution of morphological metrics relative to area increase for default parameter values (Table 1) with doubling times of *τ*_*d*_ = 9 h and *τ*_*d*_ = 20 h. The results are for one simulation run each due to very low variability between runs (see Materials and Methods for details). (a) Surface roughness *W*, (b) Coefficient of variation *CV*, (c) Relative maximum peak height *MaxP*1, and (d) Mean Fourier amplitude *I*_*k*_.

**Fig 11.**
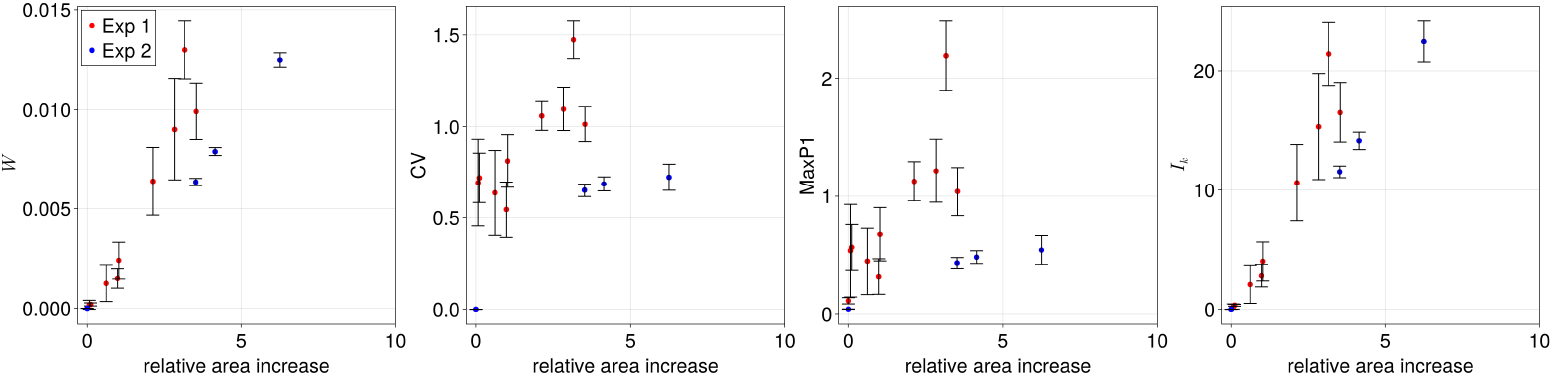
Evolution of morphological metrics relative to area increase for experimental data [47]. The dots indicate the mean and the error bars the standard deviation. (a) Surface roughness *W*, (b) Coefficient of variation *CV*, (c) Relative maximum peak height *MaxP*1, and (d) Mean Fourier amplitude *I*_*k*_.

Our results show an increase in colony growth speed and in number of fingers in simulations compared to experiments. Furthermore, we find a good agreement for the morphological metrics between the simulations and experiment 2.

### Parameter sensitivity analysis

As the baseline behaviour was established, we performed an initial testing phase with the interactive simulation mode (see Materials and Methods for details). We then systematically varied model parameters that showed the most influence on the behaviour, or are mapped directly to open questions in the field.

For all subsequent simulations, we used *τ*_*d*_ = 9 h as the default value and ran each simulation for 10 h, as this duration was sufficient for the characteristic two expansion phases to emerge (Fig. 6). We started our analysis with the agent’s movement, which is mostly determined by the orientation noise *η* and the turning rate *α* (Equation (4)).

#### Orientation noise affects colony morphology

The parameter *η* is the stochastic component of the direction alignment. The two edge cases *η* = 0 and *η* = 2*π* represent perfect chemotactic alignment and complete random direction assignment at every time step, respectively. Neither extreme represents realistic microswimmer behaviour, as their active propulsion always has a stochastic components [34]. We varied *η* in 16 steps over a range of 0.0 – 2*π* and analysed five parameter values in that range that represent the different model behaviours observed in that range (Fig. 12).

**Fig 12.**
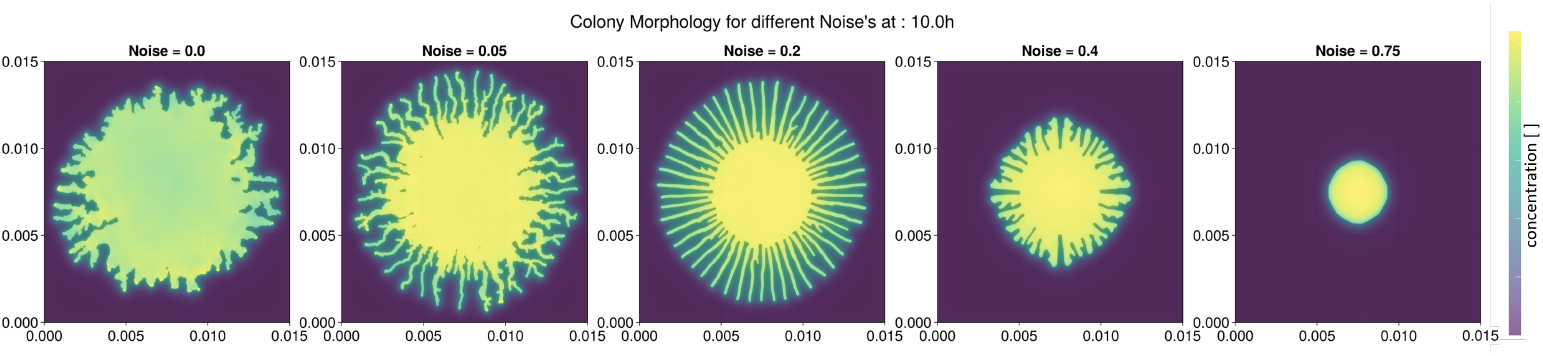
Colony morphology for five different values of the orientation noise *η* after 10 h with default parameters (Table 1). For details of visualisation see Materials and Methods.

For *η* = 0, the colony expands rapidly but irregularly, with small, irregular fingers of varying sizes. Increasing the noise slightly to *η* = 0.05 creates a more uniform gradient and more regular, finger-like expansion. At *η* = 0.2, the expansion pattern becomes much more regular, resembling the patterns observed in *in vitro* colonies. For higher noise values like *η* = 0.4, expansion is significantly slower, and fingers are less distinct. At *η* = 0.75, fingers disappear entirely, and expansion becomes exclusively circular and much slower.

The area increases faster for lower noise values (Fig. 13). Cell numbers and adsorbed material are nearly identical across all systems. The cell density increases for larger levels of noise.

**Fig 13.**
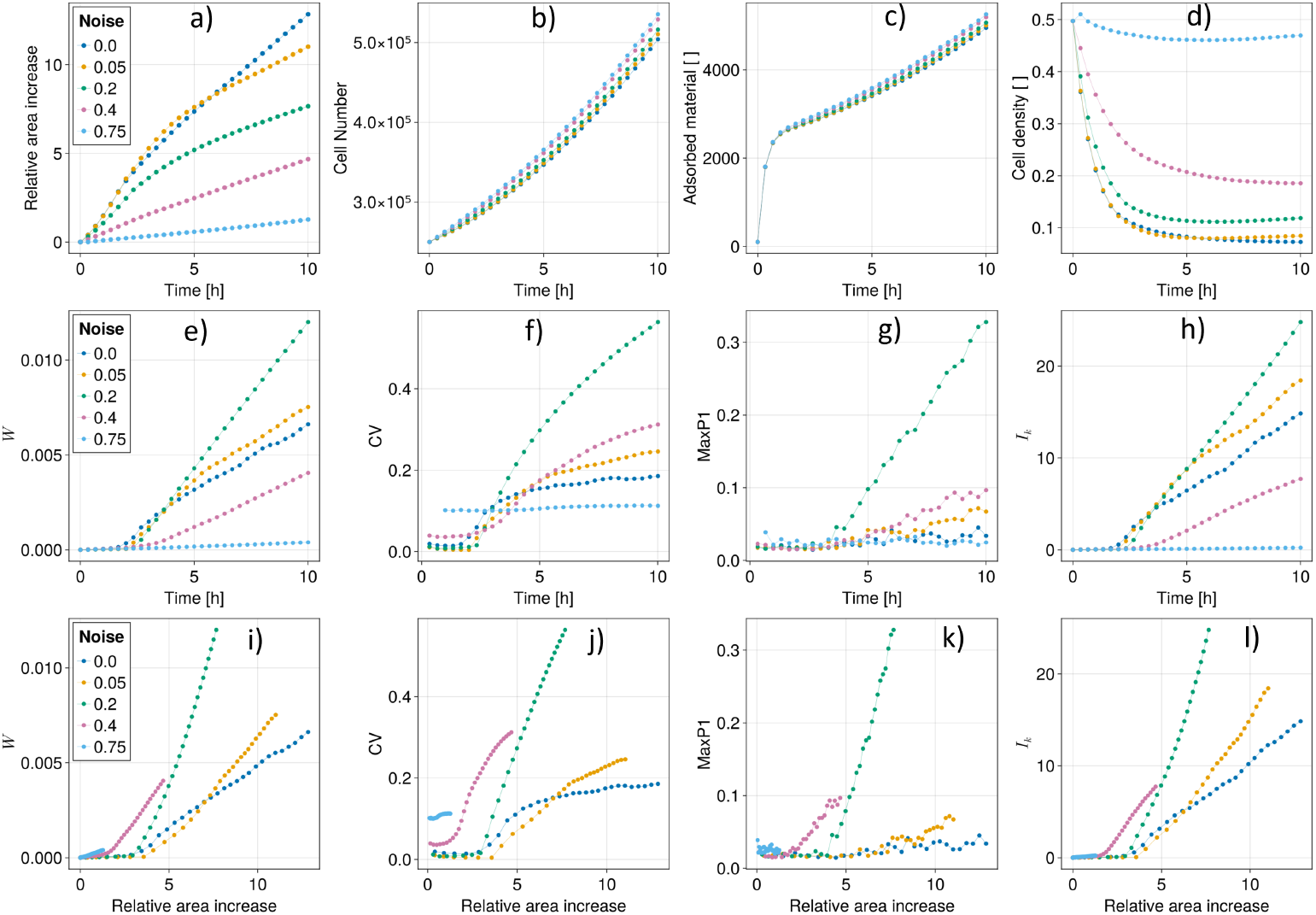
Temporal evolution of colony metrics over 10 h for default parameters (Table 1) with five different noise values in 20-minutes intervals. The results are for one simulation run each due to very low variability between runs (see Materials and Methods for details). (a) Relative area, (b) Cell number, (c) Adsorbed material, (d) Cell density within the colony, (e) Surface roughness *W*, (f) Coefficient of variation *CV*, (g) Relative maximum peak height *MaxP*1, and (h) Mean Fourier amplitude *I*_*k*_. Evolution of morphological metrics relative to area increase: (i) Surface roughness *W*, (j) Coefficient of variation *CV*, (k) Relative maximum peak height *MaxP*1, and (l) Mean Fourier amplitude *I*_*k*_.

The morphological metrics assume the highest values for orientation noise *η* = 0.2. *W* and *I*_*k*_ show bigger differences when normalized for area increase, indicating a higher sensitivity to differences in relative area increase (panels i, l). In contrast, *MaxP*1 only assumes values bigger than 0.1 for *η* = 0.2, which has very regular fingers perpendicular to the colony center. Hence, this value is highly sensitive in detecting perpendicular expansion behaviour, independent of area increase. When normalizing for relative area increase, the colony with *η* = 0.4 shows an early but less steep increase in all four metrics, indicating an early transition to finger growth but less pronounced fingers compared to the default parameters with *η* = 0.2. The *CV* shows constant values for *η* = 0.75, which is due to the slowly added, low number of grid cells to the colony, causing the grid anisotropy to become the dominant shaping factor. In summary, the orientation noise *η* has a huge impact on model behaviour, and colony development similar to *in vitro* colonies only occurs for a small range of parameter values.

#### Turning rate affects expansion speed

The turning rate *α* determines how quickly an agent responds to changes in the surrounding chemical concentration and aligns its orientation parallel to the negative gradient. The two edge cases represent instantaneous alignment (*α* = 1.0) versus no alignment (*α* = 0.0). We varied *α* in 16 steps over a range of 0.0–1.0 and analysed five parameter values in that range that represent the different model behaviours observed in that range (Fig. 14).

**Fig 14.**
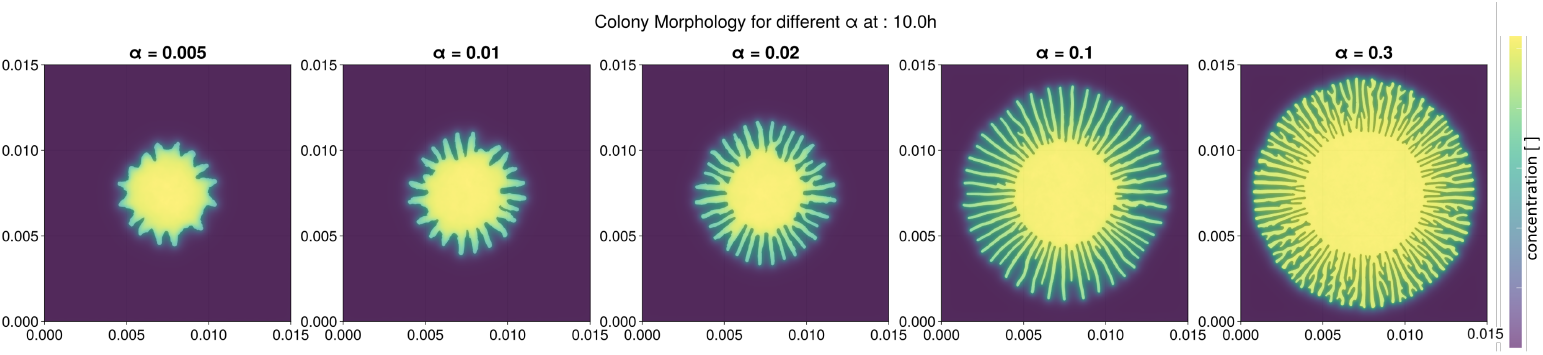
Colony morphology for five different values of the turning rate *α* after 10 h with default parameters (Table 1). For details of visualisation, see Materials and Methods.

The area increases more rapidly for higher turning rates, while cell number and adsorbed material remain nearly constant across all systems (Fig. 15). The cell density decreases as *α* is increased. Analysis of the morphological metrics reveals a complex picture. The turning rate *α* = 0.1 shows the highest values for *W* and *I*_*k*_, slightly higher than *α* = 0.3. This occurs because both metrics are sensitive to finger formation and area increase, with finger formation being stronger for *α* = 0.1 but area increase being higher for *α* = 0.3. The *CV* and *MaxP*1 metrics, which are less dependent on area increase and more sensitive to finger formation, show the highest values for *α* = 0.02, followed closely by the default value *α* = 0.1.

**Fig 15.**
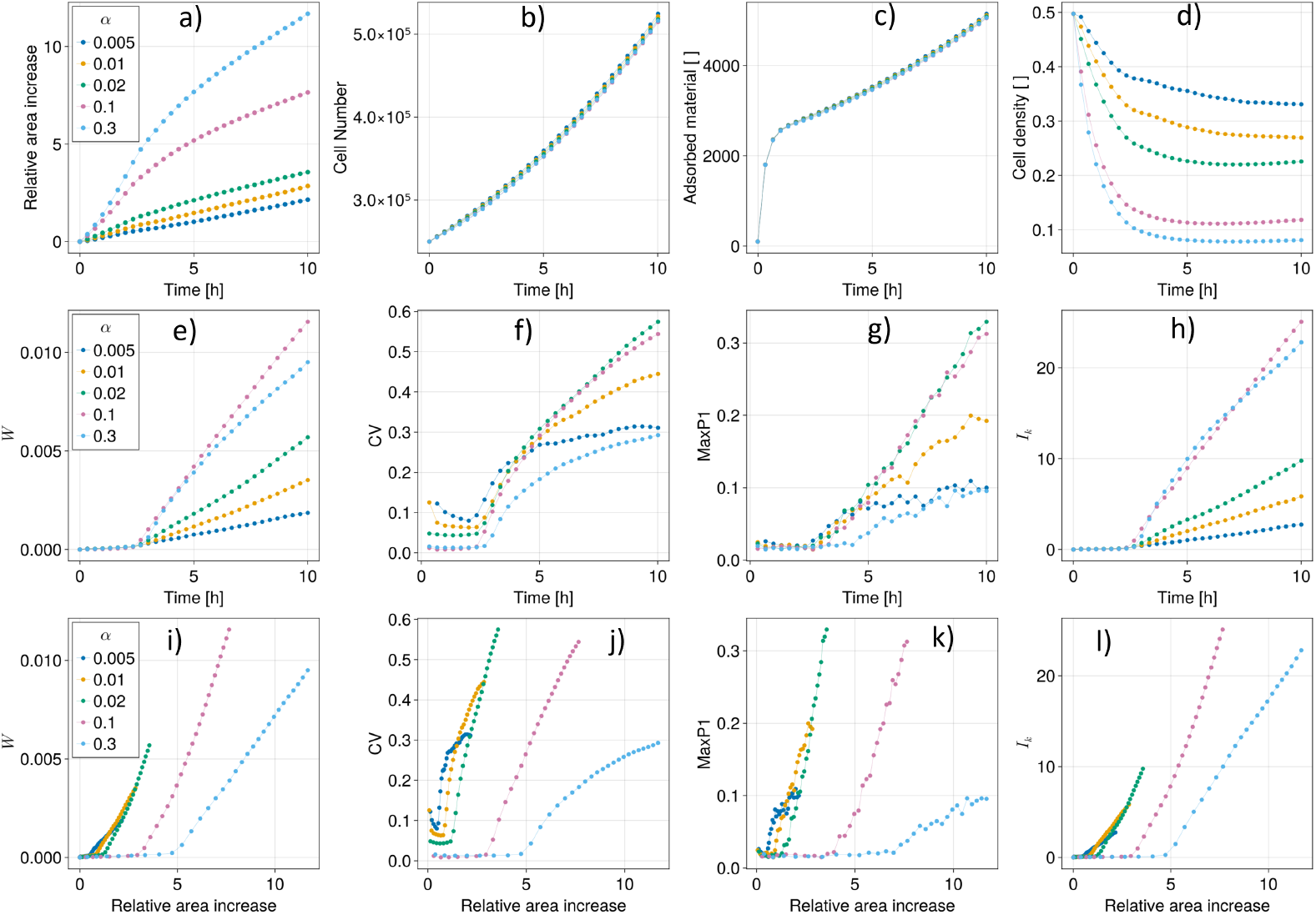
Temporal evolution of colony metrics over 10 h for default parameters (Table 1) with five different turning rates in 20-minutes intervals. The results are for one simulation run each due to very low variability between runs (see Materials and Methods for details). (a) Relative area increase, (b) Cell number, (c) Adsorbed material, (d) Cell density within the colony, (e) Surface roughness *W*, (f) Coefficient of variation *CV*, (g) Relative maximum peak height *MaxP*1, and (h) Mean Fourier amplitude *I*_*k*_. Evolution of morphological metrics relative to area increase: (i) Surface roughness *W*, (j) Coefficient of variation *CV*, (k) Relative maximum peak height *MaxP*1, and (l) Mean Fourier amplitude *I*_*k*_.

When normalized for area increase, we observe that colonies with lower values of *α* grow fingers at a much lower area value than for *α* = 0.1 and especially for *α* = 0.3. However, for very low values such as *α* = 0.005, finger formation becomes so irregular that *CV* and *MaxP*1 start to saturate after 5 h or a relative area increase of 1.

We find that the duration for the circular expansion phase is independent of the value of *α*. The value of relative area increase for the onset of fingering, however, increases with increasing turning rate *α*.

#### Small range of suitable parameter values for turning rate and orientation noise

Since both the noise *η* and the turning rate *α* influence the movement behaviour of agents, we examined their coupled effects across 20 parameter combinations that represent the different model behaviours observed in that range (Fig. 16).

**Fig 16.**
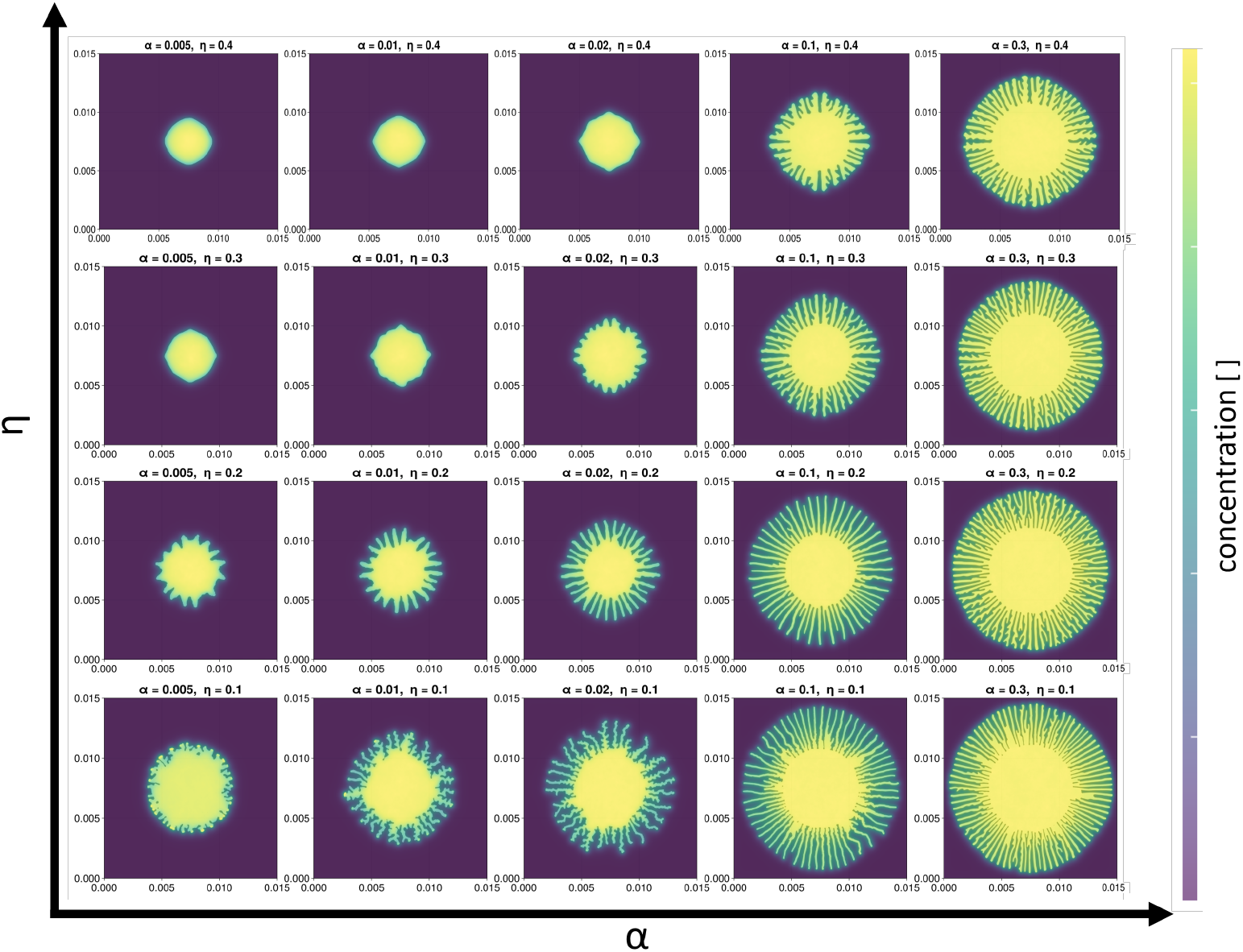
Colony morphology for five different values of the turning rate *α* (horizontal axis) and four different values of the noise *η*_0_ (vertical axis) after 10 h with otherwise default parameters (Table 1). For details of visualisation see Materials and Methods.

Within the chosen parameter range of *η* from 0.1 to 0.4 and *α* from 0.005 to 0.3, the turning rate *α* has a greater impact on colony morphology.

The noise *η* exhibits an important effect: if it is too high, lattice anisotropy becomes a defining factor in the emerging morphology. However, *η* cannot be too low either, as this leads to more irregular finger formation, especially pronounced for low values of *α*.

From visual inspection, the parameter space where expansion behaviour is similar to experiments is quite small at *η* ≈ 0.2 and *α* ≈ 0.02–0.1, a range we have already analysed (Fig. 15).

#### Grid and boundary properties affect expansion speed

After analysing the movement properties of the agents, we focused on the boundary properties in order to understand how such environmental factors impact the colony expansion. We varied the grid strength *k*_0_ from −1000 to −25000 and the grid recovery rate *g*_*rc*_ from 0.0 to 25.0 and analysed five parameter values each in those ranges that represent the different model behaviours observed (Fig. 17). To make things more intuitive to understand, we always use the absolute value of |*k*_0_| in the following sections to avoid confusion regarding the negative values for *k*_0_.

**Fig 17.**
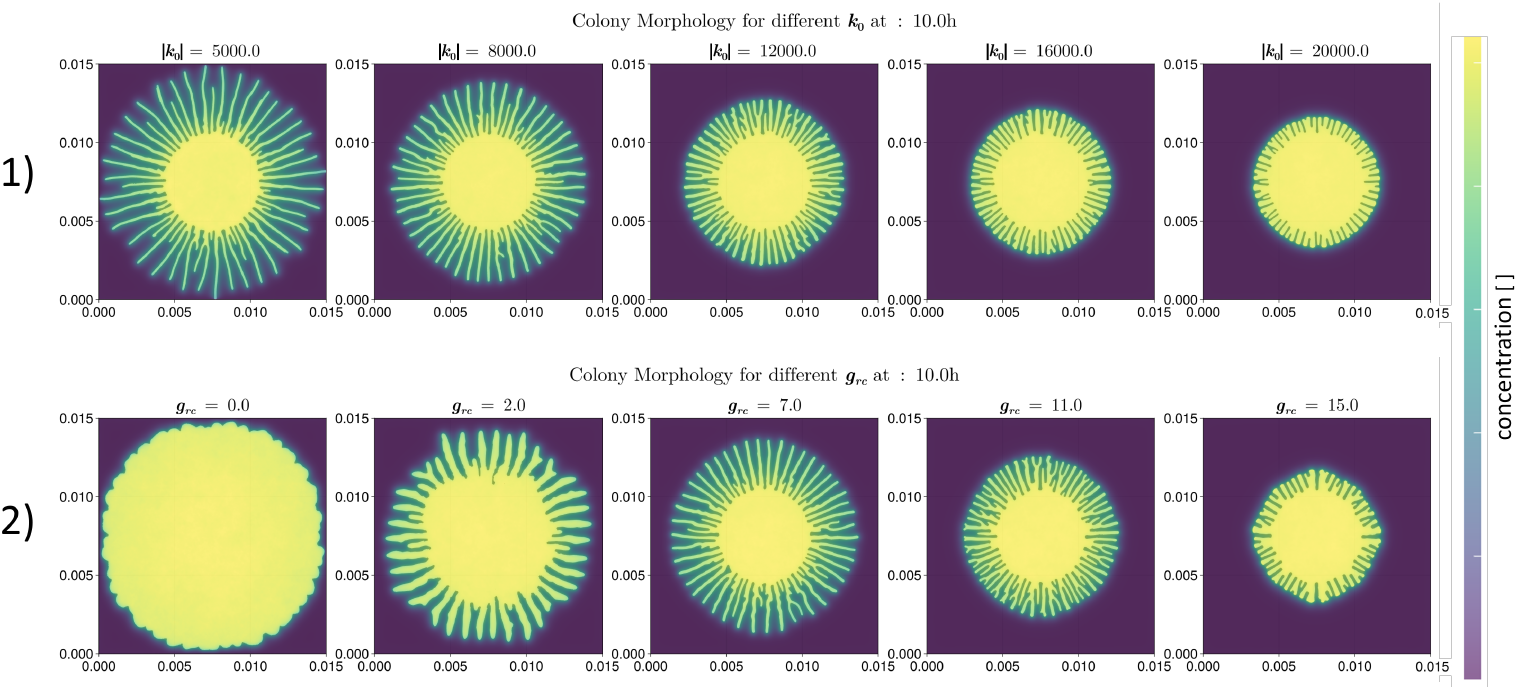
First row: Colony morphology for five different values of the grid strength |*k*_0_| after 10 h with default parameters (Table 1) Second row: Colony morphology for five different values of the grid recovery rate *g*_*rc*_ after 10 h with default parameters (Table 1). For details of visualisation see Materials and Methods.

Both a lower absolute grid strength |*k*_0_| and a lower grid recovery rate *g*_*rc*_ cause faster expansion. However, whilst a lower *g*_*rc*_ also causes the expansion behaviour to change to circular growth, a lower |*k*_0_| does not appear to change the expansion pattern.

Similarly, low values of *g*_*rc*_ of 0.0 and 2.0 show much higher relative area increase (Fig. 18), whereas all other changes in both parameters only steadily increase or decrease the relative area. Cell number and absorbed material are not shown as the behaviour is the same as for the default parameters. Cell density reflects the change in relative area.

**Fig 18.**
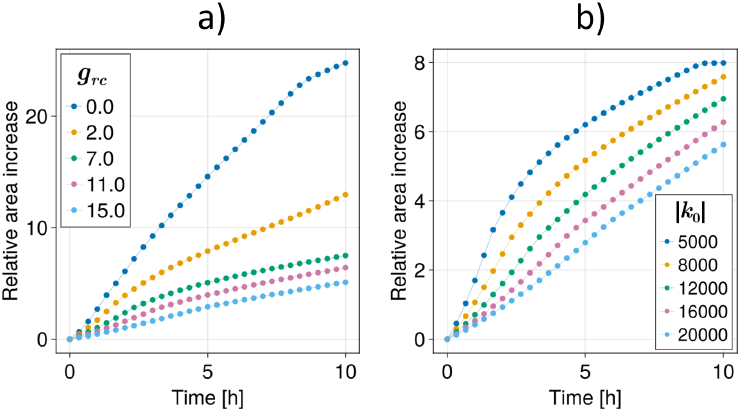
Temporal evolution of relative area increase over 10 h for default parameters (Table 1) in 20-minute intervals with (a) five different grid recovery rate values *g*_*rc*_ and (b) five different absolute grid strength values |*k*_0_ |. The results are for one simulation run each due to very low variability between runs (see Materials and Methods for details).

All morphological metrics relative to time for different values of |*k*_0_| show equidistant spacing with a later increase in values but similar slopes (Fig. 19). When normalized by relative area increase, the trend becomes clearer. Except at very high grid strength values of |*k*_0_|, all morphological metrics exhibit a marked increase within the same relative area interval of three to four.

**Fig 19.**
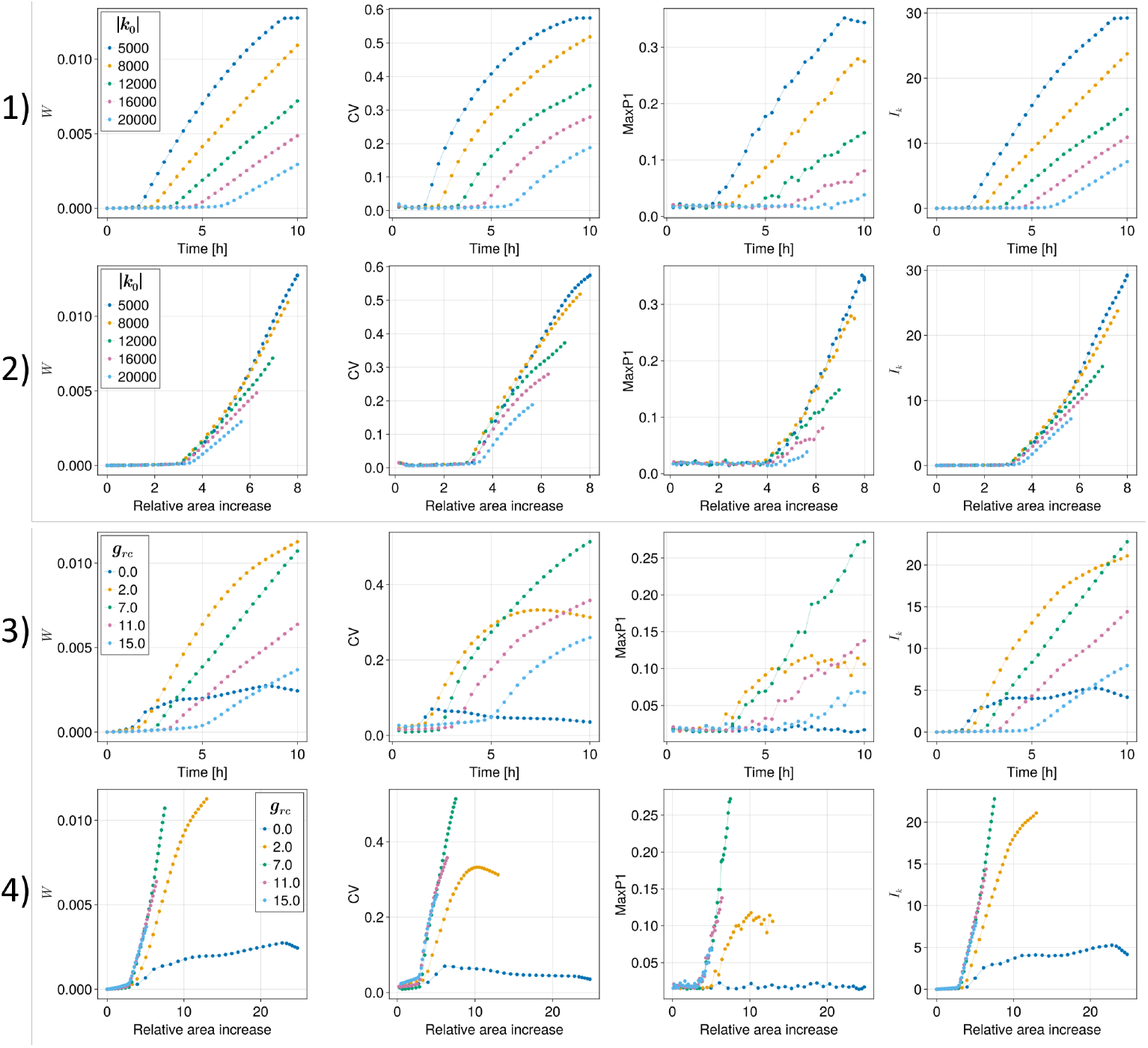
Evolution of morphological metrics over 10 h for default parameters (Table 1) in 20-minutes intervals for different grid strength values |*k*_0_| relative to time (first row) and relative to area increase (second row); for different grid recovery rate values *g*_*rc*_ relative to time (third row) and relative to area increase (fourth row). The results are for one simulation run each due to very low variability between runs (see Materials and Methods for details).

Something very similar can be observed for the grid recovery rate *g*_*rc*_, where the time point for the onset of fingering increases with increasing *g*_*rc*_. For the evolution with respect to relative area increase, all curves align except for the very low values of *g*_*rc*_ = 0.0 and *g*_*rc*_ = 2.0.

We conclude that there is a wide parameter corridor for *k*_0_ and *g*_*rc*_ that changes the temporal onset of fingering, i.e. the expansion speed but not the nature of the expansion.

#### Diffusion coefficient affects colony morphology and expansion speed

Following the previous results demonstrating that boundary properties have less influence on colony expansion behaviour than the movement properties of agents, we turned our attention to another major factor that influences agent movement: the diffusion of the chemotactically active substance. Various substances (pH/protons, glucose, cAMP, exosomes) [10–13] have been proposed as potential mediators of a chemotactic alignment in trypanosomes [11]. However, it remains unclear which specific substance could mediate the chemotactic alignment, as all are present in the experimental system.

The diffusion coefficients of these candidate substances at room temperature in water are well established in the literature, ranging from 9× 10^−9^ m^2^/s (pH/*H*^+^), 6.5× 10^−10^ m^2^/s (glucose) [51], 4× 10^−10^ m^2^/s (cAMP) [52, 53] to 7× 10^−12^–1.5× 10^−12^ m^2^/s (exosomes, depending on size) [10, 54–56]. Although the experimental setup of trypanosome colonies involves a complex liquid medium that consists of multiple components [4] placed on top of an agarose gel, previous studies have shown that the diffusion coefficients of various materials in such media differ only slightly from those in pure water [57, 58]. Therefore, we can reasonably use the diffusion coefficients measured in water as reference points.

We varied the diffusion coefficient *D*_*s*_ in our simulations across the range of biologically relevant values and studied its influence on system behaviour (Fig. 20).

**Fig 20.**
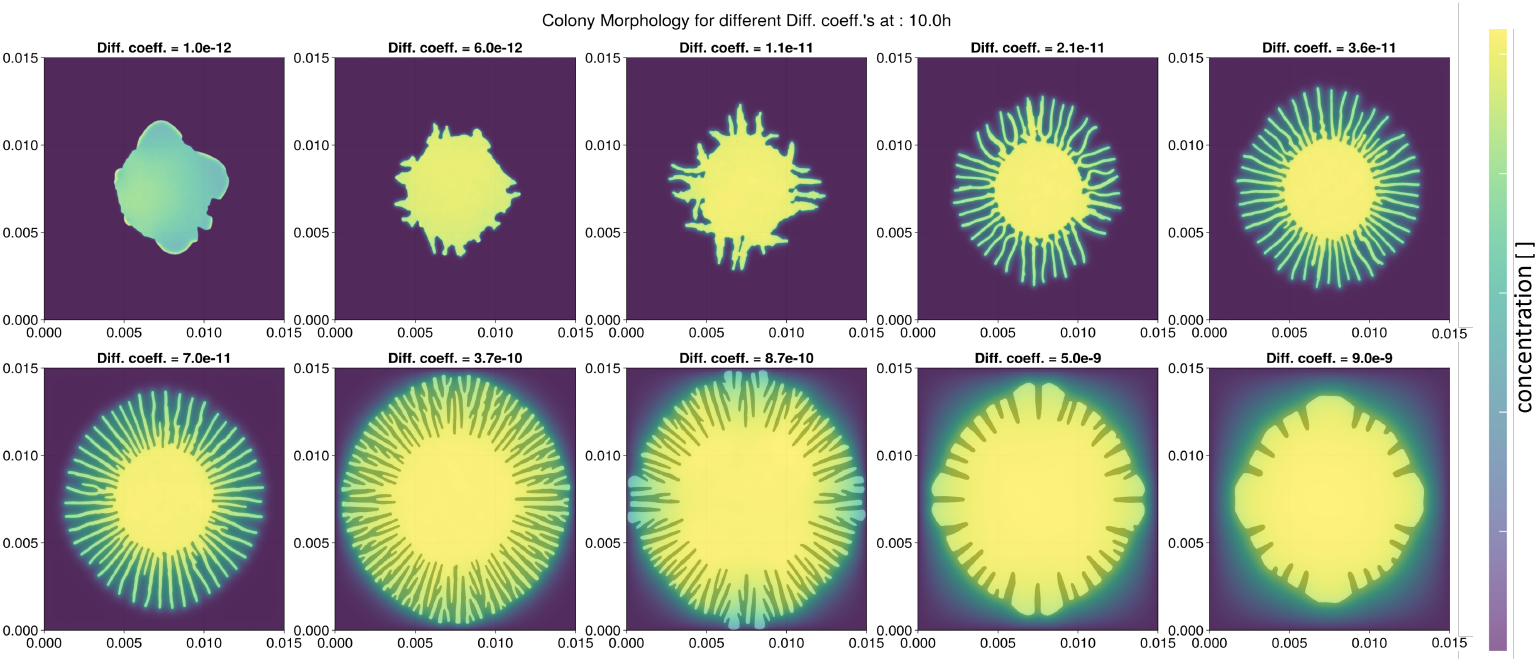
Colony morphology for ten different values of the diffusion coefficient *D*_*s*_ after 10 h with default parameters (Table 1). For details of visualisation see Materials and Methods.

The diffusion coefficient has a tremendous influence on the expansion behaviour. For the lowest value of *D*_*s*_ = 1× 10^−12^ m^2^/s, diffusion is so slow that no colony-wide concentration gradient decreasing from the center to the boundaries is established within 10 h. Instead the highest concentration is found at the boundaries of the colony. The resulting colony morphology is anisotropic and does not exhibit a finger-like pattern.

For higher diffusion coefficients, a colony-wide gradient can be established, with the highest concentration in the central part of the colony decreasing outward. This gradient becomes more and more pronounced the higher the diffusion coefficient *D*_*s*_ becomes. The resulting morphologies change steadily toward the very regular finger-like patterns observed at *D*_*s*_ = 7× 10^−11^ m^2^/s. For higher values up to 8.7× 10^−10^ m^2^/s, expansion becomes even faster, but fingers start to branch more and also merge, forming a more circular and less fractal expansion front. For even higher values of 5× 10^−9^ m^2^/s and beyond, expansion becomes slower again. Thick protrusions emerge that are not comparable to the finger-like patterns observed in experiments.

Starting at *D*_*s*_ = 3.7× 10^−10^ m^2^/s, the adsorbed chemical diffuses significantly beyond the colony fingers, which can be seen in the changing colours outside of the colony boundaries (Fig. 20), a trend that increases for larger *D*_*s*_ values. At *D*_*s*_ = 9× 10^−9^ m^2^/s, the chemical and its decreasing concentration gradient reach far beyond the colony boundaries.

The colony behaviour is the same for the first two hours (Fig. 21). After that, the area increases faster for higher diffusion coefficients until *D*_*s*_ = 8.7 *×* 10^−10^ m^2^/s. Then, the area increases more slowly for higher values of *D*_*s*_. The cell number and adsorbed material are similar for all values of *D*_*s*_ until *D*_*s*_ = 7.0 *×* 10^−11^ m^2^/s and then decrease with increasing *D*_*s*_. The cell density mirrors the relative area increase.

**Fig 21.**
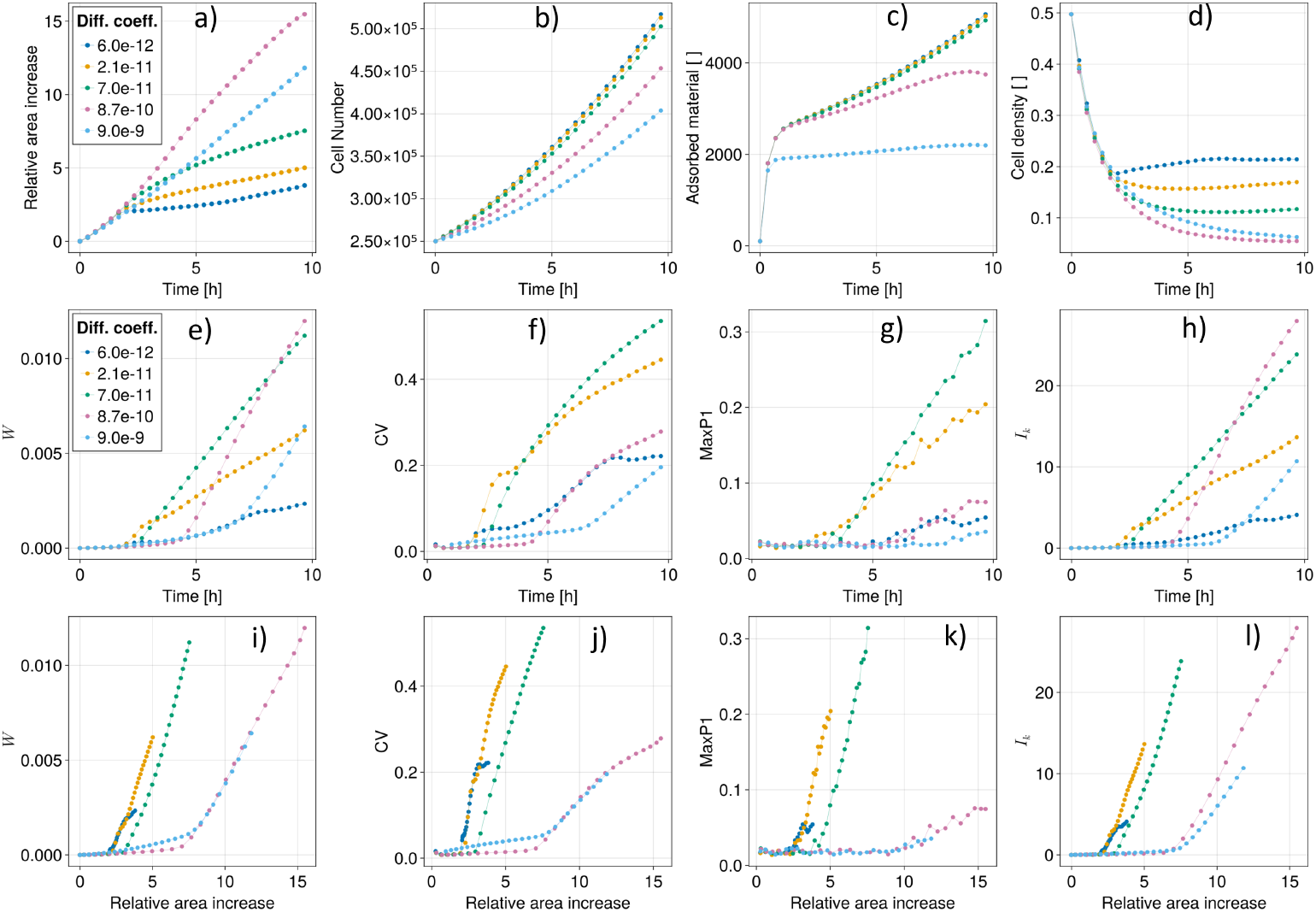
Temporal evolution of colony metrics over 10 h for default parameters (Table 1) with five representative diffusion coefficient *D*_*s*_ values in 20-minutes intervals. The results are for one simulation run each due to very low variability between runs (see Materials and Methods for details). (a) Relative area increase, (b) Cell number, (c) Total amount of material in the gradient grid, (d) Cell density within the colony, (e) Surface roughness *W*, (f) Coefficient of variation *CV*, (g) Relative maximum peak height *MaxP*1, and (h) Mean Fourier amplitude *I*_*k*_. Evolution of morphological metrics relative to area increase: (i) Surface roughness *W*, (j) Coefficient of variation *CV*, (k) Relative maximum peak height *MaxP*1, and (l) Mean Fourier amplitude *I*_*k*_. Details of visualisation in Sec..

Analysis of the morphological metrics reveals a complex picture. The default diffusion coefficient *D*_*s*_ = 7.0× 10^−11^ m^2^/s shows the highest values for *CV* and *MaxP*1, slightly higher than *D*_*s*_ = 2.1× 10^−11^ m^2^/s. This occurs because both metrics are sensitive to finger formation independent of area increase, with finger formation being slightly stronger for *D*_*s*_ = 2.1× 10^−11^ m^2^/s. This can be seen in *W* and *I*_*k*_, which are both sensitive to finger formation and area increase, showing the highest values for *D*_*s*_ = 8.7× 10^−10^ m^2^/s, slightly higher than for *D*_*s*_ = 2.1× 10^−11^ m^2^/s, as finger formation is stronger in the latter, while area increase is stronger in the former.

When normalised for area increase, all metrics behave very similarly. We observe that colonies with lower values of *D*_*s*_ start forming fingers at a lower relative area. However, for the low value of *D*_*s*_ = 6.0× 10^−12^ m^2^/s, all morphological metrics also indicate a beginning saturation in finger formation after a relative area increase of approximately 3. The system with default diffusion coefficient *D*_*s*_ = 7.0× 10^−11^ m^2^/s shows a later onset of non-circular expansion but a similar increase in the metrics, indicating similar rapid expansion in finger-like patterns. For higher values of *D*_*s*_, all metrics show a later onset of increase and a slower slope, indicating a less finger-dominated expansion pattern.

In summary, the diffusion coefficient has a clear effect on the onset of fingering with respect to relative area increase and on finger morphology.

Our parameter sensitivity analysis shows that the different parameters analysed exhibit distinct effects on model behaviour. Movement parameters *η* and *α* demonstrate a narrow parameter window for producing behaviour similar to experiments. Boundary property parameters *k*_0_ and *g*_*rc*_ primarily affect expansion speed with minimal influence on expansion pattern. The diffusion coefficient *D*_*s*_ exerts a substantial influence on expansion dynamics. Low values result in slow, irregular colony expansion. Intermediate values enhance both expansion speed and regularity. For high values, this trend reverses such that the colonies expand more slowly and regularly but exhibit progressively reduced finger formation.

## Discussion

Our agent-based model demonstrates that complex colony-level patterns in trypanosome social motility can emerge from the combination of single cell motility and proliferation, interactions of the cells with the colony boundaries and negative autochemotaxis.

Our simulation results are sensitive to the parameters for growth dynamics and single cell motility. The two simulated doubling times demonstrate a non-linear relationship between cell growth and area increase. The underlying mechanism remains unclear, though relative area growth may be primarily driven by agents at the colony surface, where density appears similar across both systems. This requires further investigation through comparison with experimental data using cell lines with different doubling times [3, 4, 12] and detailed analysis of agent position and density distributions over time within the model.

The orientation noise *η* and turning rate *α* demonstrate a narrow parameter window where finger formation is most pronounced, regular, and most similar to experiments. Such behaviour represents a remarkable result. It suggests that finger formation emerges as a direct consequence of the specific properties of the directional random walk of individual agents. Our results coincide with previous experimental findings where modifying flagellum activity and motility properties of individual trypanosomes can suppress social motility [1, 3]. Although extensive work has examined individual trypanosome movement properties both experimentally [29] and through modelling [27, 28], direct comparison with our findings remains challenging due to fundamental differences between three-dimensional free environments and our crowded two-dimensional system that constrains movement. It has been demonstrated that agent trajectories can be measured in the *in vitro* system [4, 11]. Such data is very suitable for a future direct comparison with agent trajectories extracted from the model.

The boundary parameters grid strength *k*_0_ and grid recovery rate *g*_*rc*_ exhibit a wide “parameter corridor” where changes affect only expansion speed without altering the fundamental nature of colony morphologies. This robustness suggests that the specific mechanical properties of the colony-agarose interface may be less critical than previously thought [4], with qualitative expansion behaviour being primarily determined by the interplay between chemotaxis and movement dynamics. However, our current boundary implementation is quite simplistic and could be expanded to much more complex hydrodynamic models that have been implemented for bacterial systems like Pseudomonas [23], albeit at much smaller scales or in a continuous manner where the colony boundary is treated as a free boundary problem [21, 40].

Significant computational optimisation and modification would be needed to make hydrodynamic boundary models tractable and to couple them to an agent-based model with the number of agents we have used here. Compared to our phenomenological implementation of boundaries, such models also require measurements that have not yet been done to quantify the defining physical parameter values of the hydrodynamic interface, and all of them assume some sort of lubrication caused by the microswimmers [24, 59] themselves that changes the local properties of the boundary. Whether trypanosomes secrete substances or lubricate their environment in a significant manner is currently only speculative.

The diffusion coefficient *D*_*s*_ emerges as the most influential parameter in our model, dramatically affecting both the speed and pattern of colony expansion. For low values of *D*_*s*_ = 1.0–6× 10^−12^ m^2^/s, finger formation is absent and colony growth is substantially reduced, likely because information cannot spread sufficiently rapidly across the colony to establish a global chemical gradient pointing outward toward the boundaries. Conversely, very high diffusion coefficients lead to more circular rather than finger-like growth. At these high values, signals diffuse rapidly and concentration gradients become shallow but far-reaching, extending beyond the colony boundaries. Such conditions inhibit finger formation because the highest gradient is no longer localised at the finger tips, where it guides directional growth, but instead decreases monotonically with distance from the colony centre, resulting in a unified expansion front without finger formation.

The range of simulated diffusion coefficients (10^−12^ to 10^−9^ m^2^/s) spans the known values for several proposed signalling agents, including pH (H^+^), cAMP, glucose, and small exosomes [3, 10, 12]. However, the model’s behaviour most similar to experiments occurs within the range of *D*_*s*_ = 2× 10^−11^ to 10^−10^ m^2^/s. Using the Einstein-Stokes equation:

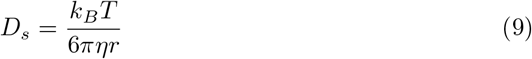

where *k*_*B*_ is the Boltzmann constant, *T* = 293.15 K is the absolute temperature, and *η* = 0.001 kg/(m·s) is the dynamic viscosity of water at *T*, the corresponding hydrodynamic radius *r* of the molecule can be estimated to be between 2–10 nm (Fig. 22).

**Fig 22.**
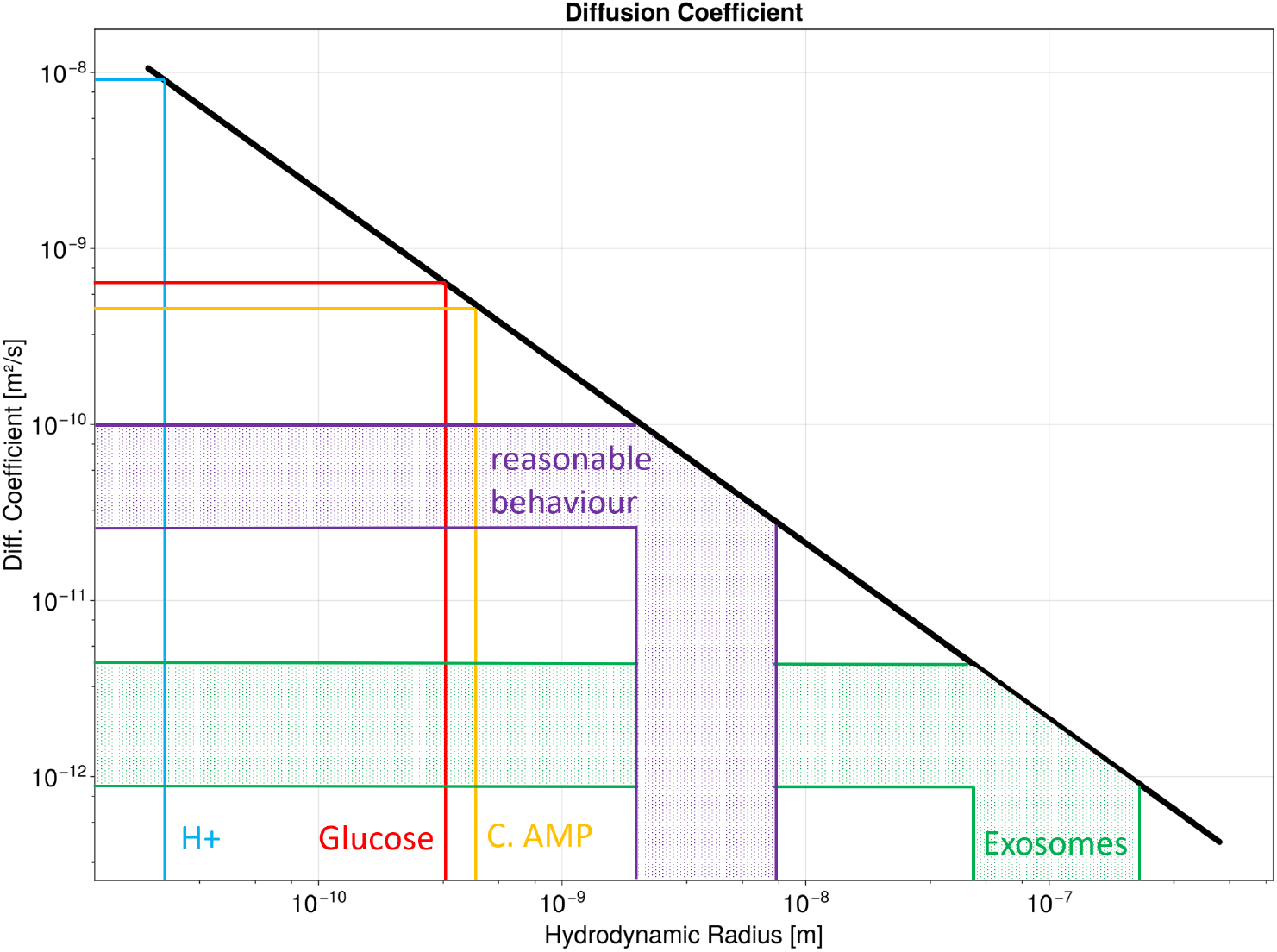
Diffusion coefficient as a function of hydrodynamic radius of diffusion particles in water according to the Einstein-Stokes equation. Model behaviour similar to experiments occurs at diffusion coefficients between 2× 10^−11^ and 10^−10^ m^2^/s, corresponding to hydrodynamic radii between 2–10 nm.

Using established conversion estimations for molecular weights [60], such radii correspond to small proteins with molecular weights between 12.1–1690 kDa. Therefore, our results suggest searching for signalling proteins/macromolecules/lipids in that weight range.

A quantitative comparison between our simulation results and experimental data shows that in the simulations, the colony growth speed and the number of fingers are larger. In contrast, the morphological metrics agree well between simulations and experiments. Hence, while the overall growth rate differs, the pattern formation dynamics remain comparable. Our parameter sensitivity analysis further indicates that adjusting the boundary strength and recovery parameters could improve the quantitative agreement.

Previous studies have shown that social motility can also occur with chiral fingers — a behaviour that is currently not captured by our model. Similarly, the inhibition of finger formation, for example, by a neighbouring E. coli colony, is not yet represented. Addressing these phenomena would require formulating hypotheses about the underlying mechanisms that could alter local motility, such as asymmetric propulsion or interspecies signalling, and subsequently incorporating these processes into the model.

## Conclusion

Our model reproduces the patterning characteristics of social motility of single colonies and demonstrates that individual movement properties of trypanosomes are critical for the emergence of finger-like patterns. Our main prediction is that boundary interactions have to be coupled with negative auto-chemotaxis and that previously proposed auto-chemotactically active substances cannot be directly responsible for social motility and likely represent correlation rather than causation. To advance this research, we have provided several testable hypotheses for experimental validation, including the motility statistics of trypanosomes, the size of potential signalling agents, and the robustness with respect to mechanical boundary properties.

## Acknowledgements

This project was funded by the Priority programme 2332 ‘Physics of Parasitism’ of the German Research Foundation (DFG).

## Supporting information

### S.1 Area of Cells

The projected two-dimensional area of trypanosomes during a social motility assay was determined using scanning electron microscope images from [4]. The cell body was manually segmented with Fiji, and the pixel count was used to calculate an approximate area of 18.0 *µ*m^2^.

**Fig S.1.**
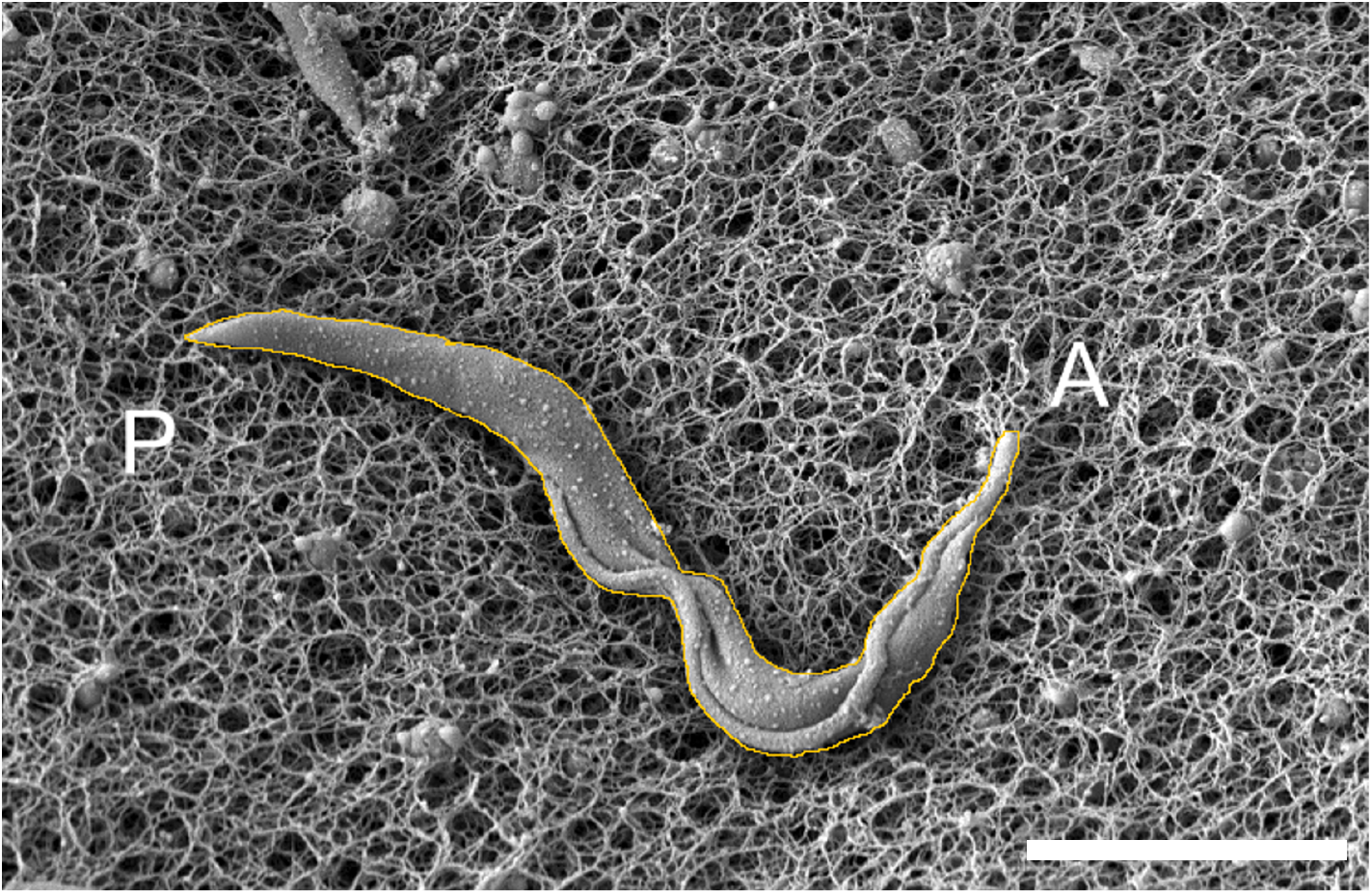
Scanning electron micrograph of a single trypanosome fixed while swimming on the surface of an agarose gel. The manually determined outline of the cell body is shown in orange. A and P indicate the anterior and posterior ends, respectively. Scale bar: 5 *µ*m. Image adapted from [4].

### S.2 Von Neumann Stability Analysis

The von Neumann stability analysis for standard diffusion equations solved with finite difference schemes has been extensively documented in the literature [44–46]. Here, we focus specifically on the stability implications when extending the analysis to our modified diffusion equation that includes additional decay and adsorption terms:

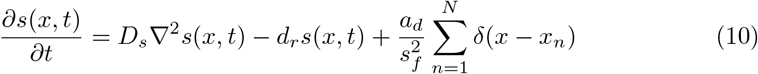

where *D*_*s*_ is the diffusion coefficient, *d*_*r*_ represents the decay rate, *s*_*f*_ is the scaling factor between the ABM grid and gradient grid, *a*_*d*_ is the adsorption rate caused by the *N* agents at their ABM grid cells *x*_*n*_, *δ* is the Dirac delta function.

For stability analysis, we consider the worst-case scenario where all ABM grid points are occupied by agents. In this case, the adsorption term 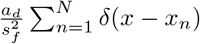 becomes a constant 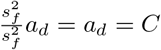 at every gradient grid point. The equation can be written in the form:

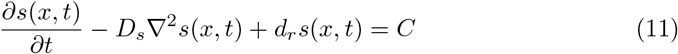

Although this equation contains a non-homogeneous term *C*, the stability of the numerical scheme depends only on the homogeneous part of the equation. This can be seen by examining the round-off error 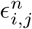, which is the difference between the numerical solution 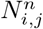 and the solution 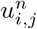 with infinite machine precision:

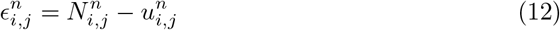

Both solutions must satisfy our discretized equation:

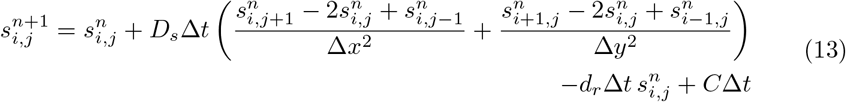

If we let *L*^∗^ represent our discretized operator, then for 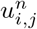 :

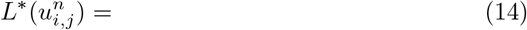

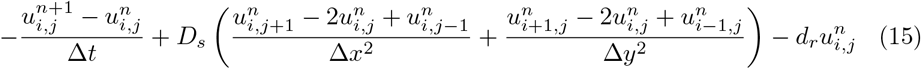

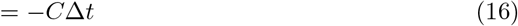

And similarly for the numerical solution:

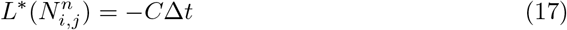

By the linearity of the operator *L*^∗^, the round-off error satisfies:

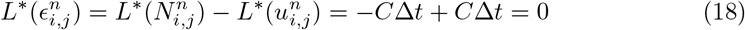

Therefore, the round-off error 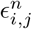 satisfies the homogeneous equation, allowing us to apply the standard von Neumann stability analysis for the explicit forward time-centered space scheme.

The derivation follows the same steps as in [45], but extended to 2D (see [61] for the general 2D case) and includes an additional decay term. Therefore, we can write 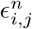 as a Fourier series:

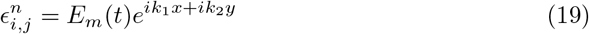

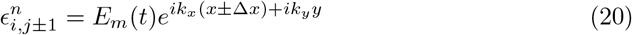

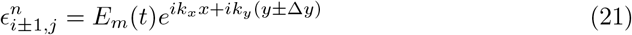

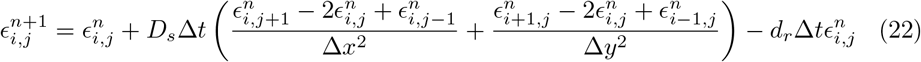

We examine the amplification factor 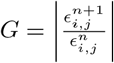 which represents the relative change of error between iterations:

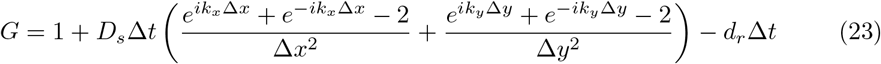

Using the identity *e*^*ik*Δ*x*^ + *e*^−*ik*Δ*x*^ = 2 cos(*k*Δ*x*), this can be rearranged to:

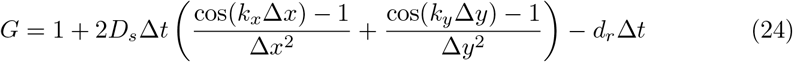

Since cos(*θ*) − 1 ≤ 0 for all *θ*, the most restrictive condition occurs when cos(*k*_*x*_Δ*x*) = cos(*k*_*y*_Δ*y*) = −1 [61], giving:

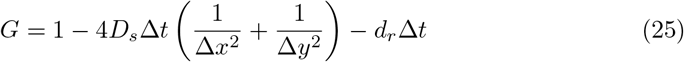

For stability, we require |*G*| ≤ 1, which leads to:

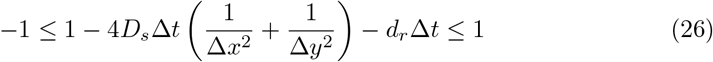

The right inequality is always satisfied, and from the left inequality we get:

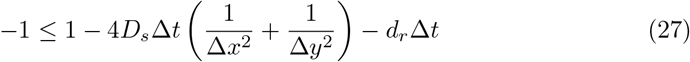

Which simplifies to:

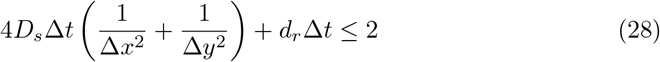

As in our simulation setup Δ*x* = Δ*y*, this can be further simplified and rearranged to our final stability condition on Δ*t*:

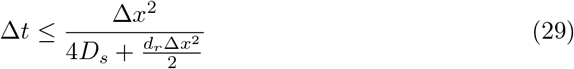

For the standard diffusion equation (when *d*_*r*_ = 0), this reduces to the familiar condition 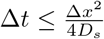 [61]. Importantly, the presence of the decay term *d*_r_ further restricts the maximum allowable time step, requiring an even smaller Δ*t* to maintain numerical stability in our simulations.

### S.3 Lattice Anisotropy

On-lattice cluster growth models can suffer from a phenomenon where the lattice symmetry becomes imprinted on the emerging structure, even with a completely random selection of growth sites [62–65]. In square lattices, this manifests as preferential growth along the major lattice axes (x and y) and their diagonals compared to other directions. This undesired anisotropy varies depending on the specific model and mitigation strategies employed. Most solutions require more complex surface site selection algorithms during the growth process to avoid anisotropic growth [63, 64, 66, 67].

Our on-lattice model of trypanosome colony geometry can be understood in a broader sense as a cluster growth model, although the colony does not “grow” in the traditional sense. Instead, swimming trypanosomes push the colony boundary outward, with each agent influencing multiple grid cells inside the circular collision area when colliding with the boundary. Nevertheless, cell growth remains a significant factor enabling colony expansion.

To investigate whether colony anisotropies do emerge in our model and how they might be mitigated, we conducted a simulation with randomly moving agents and high boundary strength (similar to the noise reduction parameter *m* in Eden-like models, which enhance growth anisotropies [62]). The simulations were normalized to produce a similar relative area increase after 10 hours.

Anisotropy emerged when using a circular collision between agents and the boundary (Fig. S.2). The colony preferentially grows along the x and y lattice axes, visible as four distinct peaks in the angular metric.

Changing the collision geometry to a diamond shape reversed this trend completely, causing the colony to grow preferentially along the diagonals of the lattice. Conversely, a square-shaped collision area dramatically increased the initially observed preferential growth along the x and y axes.

Based on these findings, we implemented a hybrid collision area combining both circular and diamond geometries. This approach effectively balanced the competing anisotropies, resulting in more isotropic colony growth that better matches experimental observations.

**Fig S.2.**
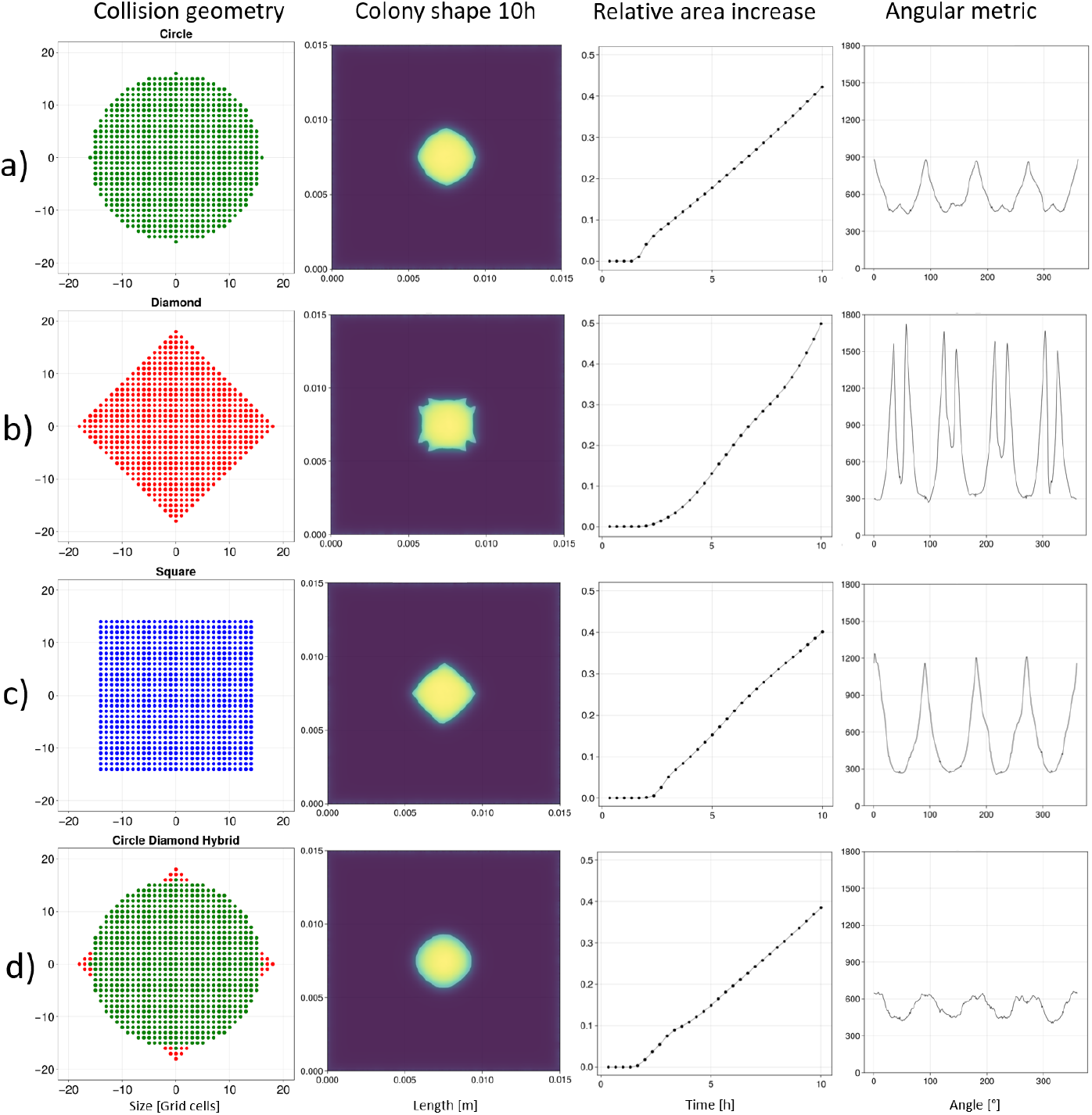
Comparison of different collision area geometries and their effect on colony growth anisotropy, normalized for similar relative area gain over 10 hours. The angular metric shows the distinct periodic anisotropies in the surface morphology. (a) Using a circular collision area results in preferential growth along the x and y axes. (b) A diamond-shaped collision area reverses this trend, favouring diagonal growth. (c) A square-shaped collision area further enhances growth along the x and y axes. (d) Hybrid collision geometry combines circle and diamond shapes to produce the most isotropic growth.

## Notes

### Competing Interest Statement

The authors have declared no competing interest.

